# Decomposing alarm-calling behaviour to investigate context-dependent threat discrimination and response escalation in a social mongoose

**DOI:** 10.64898/2026.10.16.759082

**Authors:** Zachary Cloutier, Marta B. Manser, Jack Thorley, Dominik M. Behr, Tim H. Clutton-Brock, Arpat Ozgul

**Affiliations:** Department of Evolutionary Biology and Environmental Studies, University of Zurich, Switzerland; Kalahari Research Centre, Northern Cape, South Africa; University of East Anglia, Norwich, United Kingdom; Department of Data Science and AI, Monash University, Melbourne, Australia; Department of Zoology, University of Cambridge, Cambridge, United Kingdom

**Keywords:** Ecology of fear, Cooperative breeder, Predator-prey interaction, Alarm calling, Group size, Vigilance behaviour

## Abstract

Prey can adjust their behaviour to perceived predation risk, which may induce non-consumptive fitness effects. However, quantifying perceived risk in field studies remains challenging as anti-predator behaviours often integrate multiple processes such as encounter probability and detection capacity. Partitioning prey responses into sub-processes may therefore provide novel insights on predation risk perception. To this end, we decomposed meerkat (*Suricata suricatta*) alarm-calling behaviour into threat discrimination, and response escalation. Using 23 years of data from a wild population in the Kalahari, we evaluated the context-dependence of both components in relation to landscape features, and group composition. We defined threat discrimination as the probability to emit an alarm call at predators (eagles) relative to non-predators (vultures) and response escalation as the strength and intensity of the ensuing collective reaction. We found that discrimination varied with habitat and vegetation productivity. Furthermore, larger groups and the presence of pups increased meerkat threat discrimination, but only during the dry season. Response escalation varied with habitat and group size, but pup presence only increased reaction strength. The contrasting context-dependence among the two alarm-calling components support decomposing compound behaviours rather than quantifying perceived risk from a single behaviour. Thus, we recommend future studies to use this approach, when possible, to improve inference on prey perception and responses to risk.

## Introduction

By shaping behavioural and physiological responses to the risk of mortality, predation is a critical driver of prey ecology and evolution (Lima and Dill 1990; Brown et al. 1999; Prugh et al. 2019). When directly attacked, prey rely on reactive responses to predators such as flight (Sunquist and Sunquist 1989; Creel 2018). Under sustained high predation risk, however, prey also engage in proactive anti-predator behaviours including vigilance and avoidance that may improve short-term survival (Boonstra 2013; Creel 2018). These responses are not without costs: anti-predator tactics can require substantial time and energy, generating non-consumptive effects on fitness (Lima 1998; Zanette et al. 2011; Creel 2018; Prugh et al. 2019; Peacor et al. 2020; Kelleher et al. 2021). For example, intense predation pressure from cougars (*Puma concolor*) indirectly reduced bighorn sheep (O*vis canadensis)* growth, reproductive output, and neonatal survival (Cloutier et al. 2024).

Despite recognition that the accurate assessment of predation risk is important for prey fitness (Stankowich and Blumstein 2005; Cresswell 2008), the quantification of perceived risk in natural environments remains challenging (Moll et al. 2017; West et al. 2018; Prugh et al. 2019; Driscoll et al. 2022). Most studies aiming to do so rely on encounter-based metrics of predators and prey, but these may not reflect the risk perceived by prey as predator presence does not necessarily imply an active hunting behavioural state (Abrahms et al. 2016; Prugh et al. 2019). Because non-consumptive effects arise from prey response to *perceived* rather than *actual* predation risk (Cresswell 2008; Moll et al. 2017; Creel 2018), behavioural responses to predators can offer useful indicators of risk perception; such as giving-up densities, vigilance rates, or flight initiation distances (Brown 1988; Camp et al. 2012; Wagnon et al. 2020).

Alarm signals are immediate proactive response to detected threats (McLachlan and Magrath 2020; McDonald et al. 2022) and can initiate coordinated reactions in group-living animals (Bradbury and Vehrencamp 2011; Kappeler 2021). Both the propensity to produce alarm calls and the strength of the subsequent response are expected to vary with perceived risk (Lima and Dill 1990). While alarm calling can be relatively simple in some species, several species evolved alarm call systems that convey detailed information about the type, location, or magnitude of threat (McLachlan and Magrath 2020; McDonald et al. 2022; Yu et al. 2025). Vervet monkeys (*Cercopithecus aethiops*) are a classic example as they emit predator-specific alarm calls in response to leopards (*Panthera pardus*), martial eagles (*Polemaetus bellicosus*), and pythons (*Python sebae*; Seyfarth et al. 1980). Functionally referential predator calls have also been demonstrated in Diana monkeys (*Cercopithecus diana diana*; Zuberbühler et al. 1997), black-fronted titi monkeys (*Callicebus nigrifrons*; Cäsar et al. 2012), saddleback and moustached tamarins (S*aguinus fuscicollis* and Saguinus mystax; Kirchhof and Hammerschmidt 2006), as well as Gunnison’s prairie dogs (*Cynomys gunnisoni*; Placer and Slobodchikoff 2000).

Here, we use the functionally referential alarm calling system of meerkats (*Suricata suricatta*) to investigate how predation risk is detected, assessed, and acted upon. Meerkats are small, diurnal, highly social carnivores that are exposed to a diverse range of terrestrial and aerial predators (Clutton-Brock et al. 1999a). Their alarm calls have been shown to vary according to predator type and perceived urgency, allowing efficient collective reactions (Tables S1.1 and S1.2; Manser 2001; Manser et al. 2002, 2008). Following an alarm call, meerkats can exhibit a weak response by briefly looking at the threat, or strong responses by interrupting foraging to flee to refugia (Manser and Bell 2004), mob terrestrial predators (Graw and Manser 2007), or engage in bipedal vigilance (Clutton-Brock et al. 1999c). Reaction strength can therefore provide information about the urgency perceived by the signaller (Manser et al. 2002). Alarm calls are often emitted by dedicated sentinels who temporarily forgo foraging to scan for potential threats, allowing other group members to concentrate on foraging (Rauber and Manser 2017).

Interpreting alarm calling as an indicator of perceived predation risk requires recognising that it is a compound behavioural process. Alarm call emission reflects the occurrence and detection of potential threats as well as the signaller’s assessment of their risk, while the subsequent collective response provides further information about how that stimulus is perceived. Although the meerkat alarm-calling system has been well studied (Manser 1999; Manser et al. 2002; Hollén and Manser 2007; Schibler and Manser 2007; Townsend et al. 2012), it is unclear whether the environmental and social contexts influencing alarm-calling propensity are the same as those shaping threat discrimination and the strength of subsequent collective responses. Recorded alarms also depend on the probability that an event is observed and recorded, so recorded alarm occurrence cannot be read directly as perceived risk. Conditioning on a recorded alarm reduces this problem: comparing alarms to predators and non-predators removes nuisance variation in total encounter, detection, and observation processes insofar as these affect both stimulus types similarly, although species-specific differences in availability and detectability remain. Analysing the group’s response following a recorded non-predator alarm then examines how strongly meerkats respond once such a stimulus has been detected and called at, although characteristics of the encounter, such as the bird’s distance or altitude, may still vary.

In this study, we investigated predation risk perception by decomposing alarm calling behaviour into sequential processes using 23 years of behavioural data from a wild meerkat population in the Kalahari: (1) alarm-calling propensity, (2) threat discrimination, and (3) escalation of the subsequent collective response (Figure 1). This framework allows us to separate variation in encounter and detection of threats from variation in threat discrimination and perceived urgency. To isolate drivers of alarm calling representative of meerkat’s perspective, we used alarm calls in response to soaring predators (eagles) and non-predators (vultures), which roam over wide ranges at high altitudes and can thereby be detected over long distances. Across all analyses, we accounted for the influence of spatial, weather, and social contexts on encounter probability, detection capacity, threat level, and perceived risk. These contexts can affect each step differently: habitat structure, vegetation, and topography can alter vulnerability and sightlines (Lagory 1986; van der Merwe and Brown 2008; Embar et al. 2011; Camp et al. 2012; White et al. 2022; Lennox et al. 2025), weather can affect behaviour, detection, and communication (Hayes and Huntly 2005; Beauchamp 2010; Camp et al. 2012; Cherry and Barton 2017; Rabaiotti and Woodroffe 2019), and group composition can change both detection capacity and vulnerability (Pulliam 1973; Lima 1995; Lea and Blumstein 2011; Lehtonen and Jaatinen 2016). Because alarm-calling propensity is itself a compound outcome, we describe it only as environmental context (Appendix S1) and base our inference on two questions, both conditional on a recorded alarm to a high-soaring bird. First, does context affect predator–non-predator discrimination, indexed by the probability that an alarm was directed at a predator (eagle) rather than a non-predator (vulture)? Second, following non-predator alarms, does context affect escalation of the ensuing collective reaction, first from a weak to a strong reaction and then from bipedal vigilance to fleeing (Figure 1)? By analysing alarm-calling propensity, threat discrimination, and response escalation as successive stages, we decomposed meerkat anti-predator behaviour rather than collapsing it into a single process.

**Figure 1.**
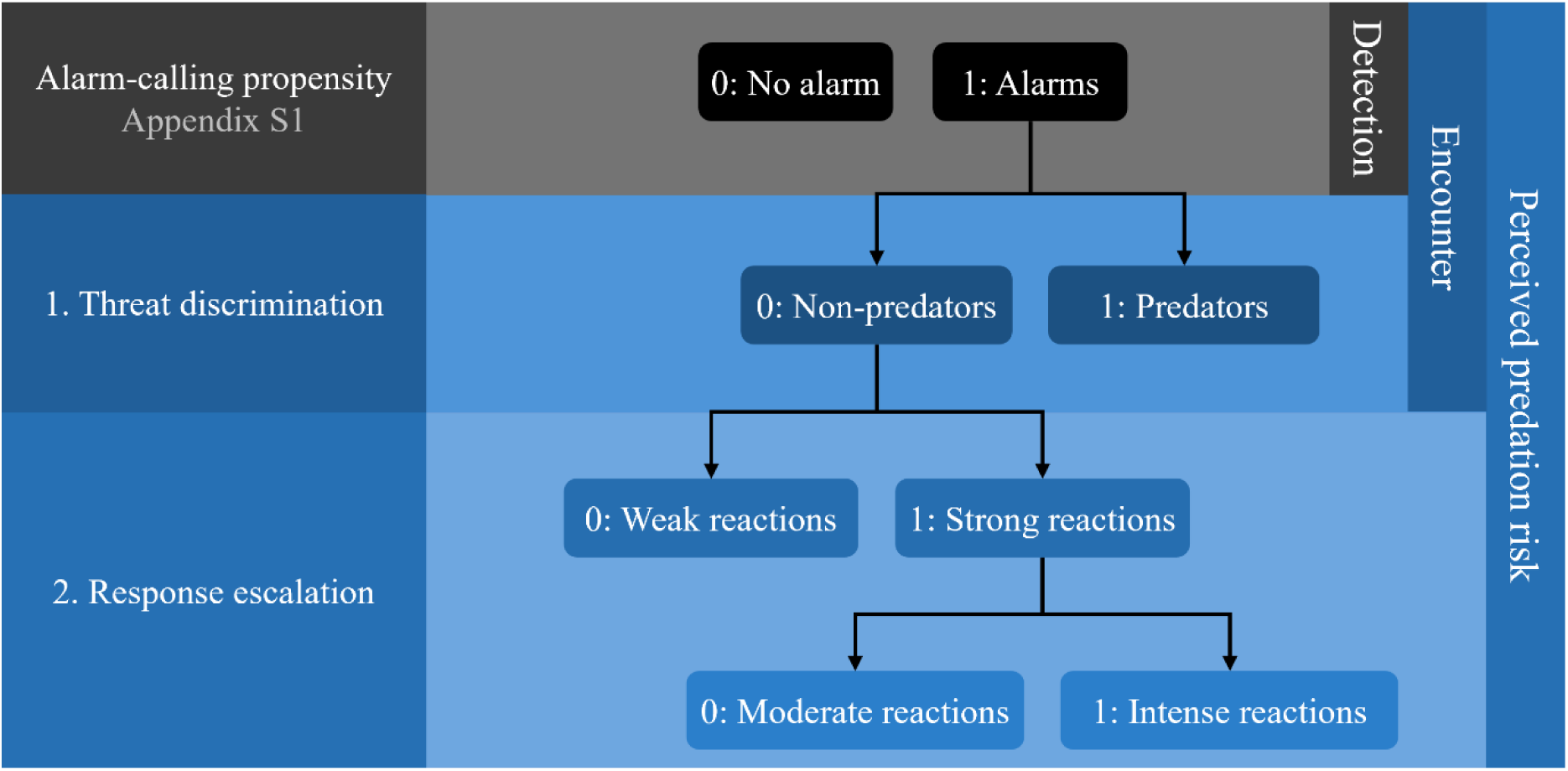
Scheme of the sequential framework. Recorded alarm occurrence is a compound outcome of encounter, detection, the decision to call, and observation, and is described only as context (Appendix S1). The main analyses are conditional on a recorded alarm to a high-soaring bird: predator–non-predator discrimination, and escalation of the collective response following non-predator alarms.

## Methods

### Study Area

The study population is located at the Kalahari Research Centre (KRC; 26°58’S, 21°49’E), in the Northern Cape province of South Africa (Clutton-Brock and Manser 2016). The 73.5 km^2^ study site comprises heterogeneous habitats across an altitudinal range of 901 m to 961 m, and is characterised by a fossil riverbed running through a duneveld landscape (Thorley et al. 2026). The site is surrounded by livestock farms and game reserves and is enclosed by a fence that keeps out large terrestrial species without limiting the movements of meerkats and other smaller species. Therefore, the study area contains fewer terrestrial predators than the wider Kalahari environment (Clutton-Brock et al. 1999a).

The climate is semi-arid and characterized by a warm, wet season from October to April, followed by a cold, dry season (Russell et al. 2002; Maag et al. 2019), which overlaps with the breeding season of both eagles and vultures (Maclean 1969; Ferguson-Lees 2005). During the wet season, daily maximum temperatures average 34.3°C (± 4.6°C), ranging from 14.1°C to 44.4°C. In contrast, the dry season average drops to 25.5°C (± 5.5°C), with maximum daily temperatures varying from 2.0°C to 39.9°C. Coupled with rainfall variations, these seasonally dynamic conditions drastically change the vegetation landscape and food availability (Thorley et al. 2025a, b). As a result, meerkat litter production is higher during the wet season (Clutton-Brock et al. 1999b).

### Meerkat Monitoring and Alarm Call Data

Each habituated meerkat group was followed in the morning and afternoon for an average of 3-4 days per week (Clutton-Brock and Manser 2016). During these follows, alarm calls were recorded *ad libitum*. Meerkats emit distinct predator-specific alarm calls when they perceive a threatening presence of birds, mammals or snakes/secondary predator cues (Manser 2001). They will further emit three other alarm call types (alert, panic, and moving animal calls) that are not predator-specific (Manser 2001). ‘Alert’ calls are produced in response to aerial non-predatory birds at proximity or raptors (predators or not) that are more than 500 m away and usually generate a weak collective response (Manser 2001). ‘Panic’ calls are produced in response to another species’ alarm call or when any type of predator makes a sudden movement, generally generating a strong response by fleeing to a bolthole (Manser 2001). Finally, ‘moving animal’ calls are reactions to distant movement from any animal type (Manser 2001). Acoustic analyses revealed differences between alarm calls of low, medium, and high urgency independently of the stimulus triggering them (Manser et al. 2002).

Previous work found that most recorded alarm calls (92.6%) were in response to aerial rather than terrestrial threats (Thorley and Clutton-Brock 2017). Because alarm calls can be elicited by a wide range of both biotic and abiotic stimuli that vary in their spatial and temporal distributions, we restricted analyses to responses to high-soaring birds. These birds roam over wide areas and can be detected by meerkats over long distances, making their encounter rate less dependent on local habitat and allowing the location of an alarm call to more closely represent the spatial context experienced by the meerkats. Furthermore, meerkats can detect them over long distances as they soar at high altitudes. High-soaring birds we classified as predators (eagles) or non-predators (vultures; see Tables S1.1 and S1.2).

Human observers recorded alarm calls, which are conspicuous and usually spatially concentrated, only when at least 50% of the group reacted and documented the ensuing collective reaction from the majority of the group. We considered ‘briefly looking at the stimuli’ as a weak response, whereas bipedal vigilance and fleeing were categorized as strong reactions as meerkats cease foraging. We further separated strong reactions by classifying bipedal vigilance as moderate and fleeing as intense. We excluded ‘mobbing’ from our analyses.

Between 2002 and 2024, GPS locations were recorded approximately every 15 minutes during group ‘follows’ using a hand-held GPS (Garmin eTrex 10; Garmin ltd) from the centre of the group (Kranstauber et al. 2020). Sleeping burrows were also recorded for the same period, while boltholes were only recorded sporadically between 2002 and 2007, then continuously from 2022. Each alarm call was matched to the closest GPS location within a 15-minute window. The mean lag time between an alarm call and its associated location was 4:06 ± 2:44 mins (mean ± SD).

### Spatial, Weather, and Social Variables

To characterize the heterogeneity of the landscape, we used habitat type and Normalized Difference Vegetation Index (NDVI). These spatial variables capture the structural and vegetative properties of the environment related to meerkat risk perception as they reflect spatial variation in refuge availability, vegetation cover, and detection capacity (Bateman et al. 2015; Thorley and Clutton-Brock 2017; Thorley et al. 2026). Habitat classification followed established descriptions of the region from van Rooyen et al. (2008), accounting for soil characteristics and the vegetation community. The vegetation is dominated by perennial grasses including *Eragrostis*, *Aristida*, *Stipagrostis*, and *Schmidtia*, alongside dwarf shrubs and tree cover (*Vachellia sp.* and *Boscia* in particular). Four habitat categories were identified at the study site from a Landsat-based vegetation map (Figure 2; Thorley et al. 2026), listed here in order of increasing structural complexity: riverbed and pans, high duneveld, low duneveld, and low shrublands (see Table S1.3 for descriptions).

**Figure 2.**
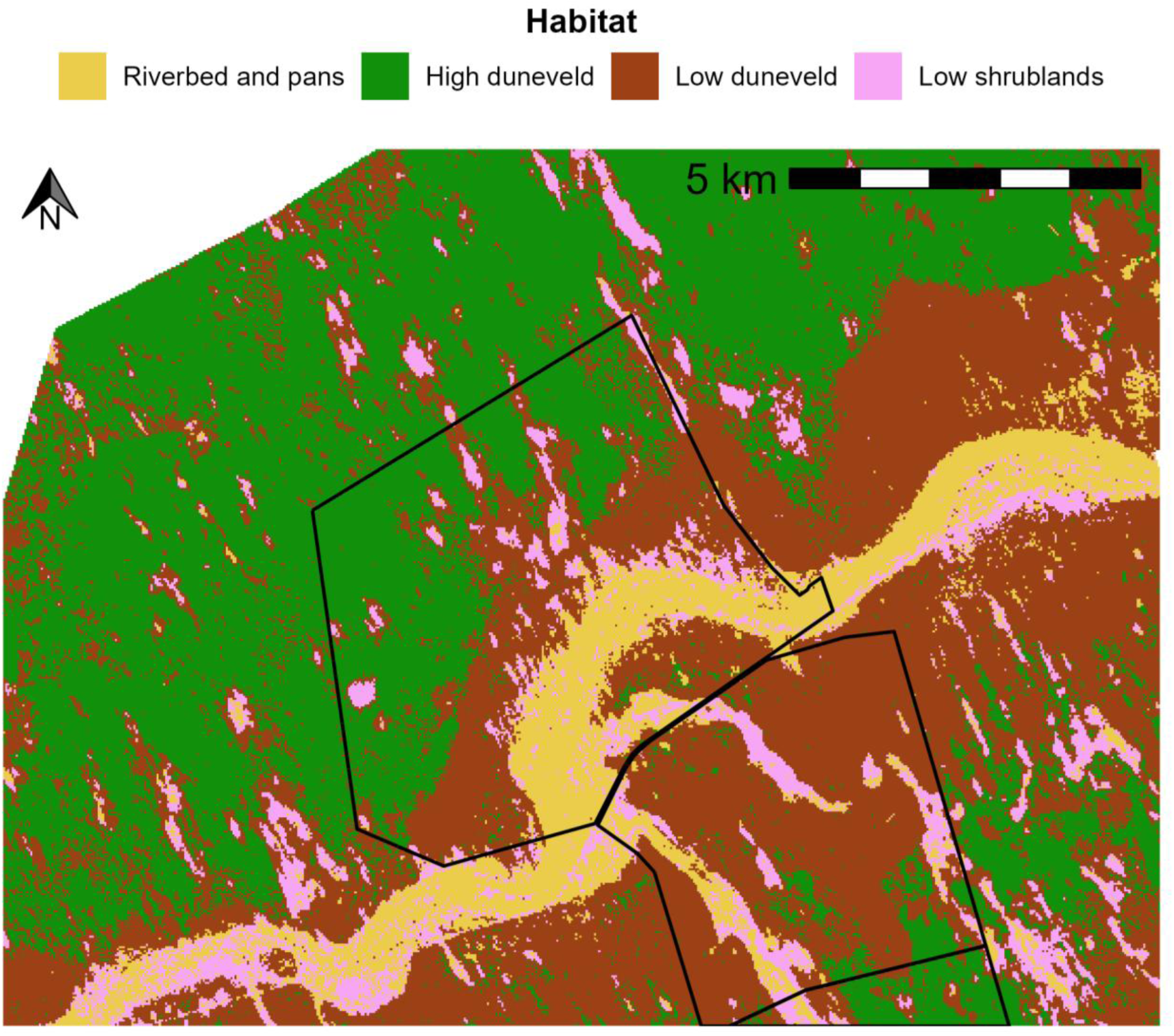
Habitat map of the Kalahari Research Centre, South Africa. Habitats were classified in four categories: riverbed and pans, high duneveld, low duneveld, and low shrublands. The black borders show the fenced perimeter of the main farms within the study area.

To assign habitat to each location while accounting for group spread, we extracted the most dominant habitat type within a 55-m buffer, corresponding to the average distance covered by a group in 15 minutes of morning foraging (54.5 m, calculated from a mean hourly speed of 217.5 m ± 193.8 m; Thorley et al. 2026). We extracted NDVI, which captures vegetation productivity, for every 16 day-windows between 2002 and 2024 at a 250-m resolution from MODIS MOD13Q1 product (Didan 2015). Habitat type and NDVI were only weakly correlated (maximum point-biserial correlation: -0.09), so both variables were retained.

Weather conditions during each follow were recorded as either fine, windy, overcast, rainy, or any combination thereof. From this, we created a binary visibility variable comparing overcast conditions (1) likely to impair visibility and therefore predator detection, to fine conditions (0). We further created another binary variable for acoustic conditions comparing only windy conditions (1), which may affect acoustic transmission, to fine conditions (0).

We included group size and the presence of pups as social predictors. Group composition was recorded by observers at the start of each follow (Clutton-Brock and Manser 2016). We classed group size as the daily median number of individuals sighted within a group. The presence (1) or absence (0) of pups foraging with the group was recorded as a binary variable.

### Sample Size

The final dataset for the wet season consisted of 13,803 alarm calls, for which 3,647 were in response to predators and 10,156 to non-predators. For the dry season, the dataset consisted of 14,774 alarm calls, 2,806 directed at predators and 11,968 at non-predators. Across both seasons, alarm calls to non-predators elicited 4,495 weak and 17,103 strong collective responses. Within strong responses, 6,273 were classified as moderate and 10,790 as intense.

### Statistical Analyses

We analysed the two conditional components of meerkat alarm calling separately (Figure 1). We fitted models using the *glmmTMB* package version 1.1.13 (Brooks et al. 2017). All interactions and their associated biological rationale are listed in Table S1.4. Numerical predictors were standardized using z-score transformation (mean = 0, SD = 1) to facilitate effect size comparison. To ensure convergence, we used the *nlminb* optimizer and increased iteration limits to 1,000. All models converged without warnings. We describe the spatial distribution of refugia and the environmental context of alarm-calling propensity in Appendix S1. Because alarm-calling propensity is a compound outcome, we did not use it as a measure of perceived risk.

We then evaluated predator–non-predator discrimination by comparing alarm calls made in response to predators (1) to those made in response to non-predators (0) in each season by fitting GLMMs with a binomial error structure. The dataset consisted of all recorded alarm calls to eagles and vultures. Conditioning on a recorded alarm to a high-soaring bird removes nuisance variation in total encounter, detection, and observation processes insofar as these affect eagle and vulture alarms similarly. It does not remove species-specific differences in the relative abundance, detectability, or identifiability of eagles and vultures, which are therefore the central assumption of this analysis: context effects on this response can be interpreted as differences in discrimination only if such variation does not covary importantly with the predictor of interest, is controlled for, or can be shown not to explain the result. Season-specific models absorb the main breeding cycles of both groups and the year random intercept absorbs between-year differences in their average relative abundance. Recorded alarm calls are conditional on at least half of the group reacting, which could be affected by group size. Because relative abundance may also change within seasons, the interpretation of context effects that covary with within-season timing, particularly pup presence, is conditional on a within-season sensitivity analysis. We repeated the threat-discrimination analysis after excluding riverbeds and pans, which contain most of the large trees that provide nesting and perching sites for large birds of prey and vultures. This sensitivity analysis tested whether our results were robust to the inclusion of habitats where encounters may more frequently involve perched birds rather than birds soaring overhead. As predictors representing meerkats’ vulnerability and uncertainty, we considered habitat type and NDVI for spatial context, visibility conditions for weather context, as well as group size and pup presence for social context. We included a two-way interaction term for habitat type and NDVI. We accounted for pseudoreplication by fitting group identity and year as random intercepts.

We analysed meerkat response escalation using alarm calls to non-predators only. Responses to predators were almost always strong (97% of eagle alarm calls with an assigned reaction, compared with 79% for vultures; Table S1.2), so predator identity would dominate any pooled analysis. Restricting the analysis to non-predators therefore allowed us to examine contextual variation in response thresholds towards a stimulus that is not itself a direct threat, which captures one component of perceived risk. To do so, we fitted binomial GLMMs comparing strong (1: bipedal vigilance and fleeing) to weak (0: briefly looking at the stimuli) collective reaction to alarm calls. We then further disentangled strong reactions, by comparing intense (1: fleeing) to moderate (0: bipedal vigilance) collective reactions. Together, these two binary models form a sequential (continuation-ratio) ordinal model of response escalation, which allows contextual effects to differ between the transition from weak to strong reactions and the transition from vigilance to fleeing. For both analyses, we included habitat type, acoustic conditions, group size and pups as predictor for spatial, weather, and social contexts. Consistent with the threat discrimination analysis, we fitted group identity and year as random intercepts to account for pseudoreplication.

For each analysis, we built 11 candidate models according to the four *a priori* predictions for spatial, weather, and social contexts, as well as habitat-specific context-dependence. We then used the package *AICcmodavg* version 2.3.4 (Mazerolle 2023) to compare candidate models with the corrected Akaike information criterion (AIC_c_). In cases where ΔAIC_c_ < 2, we followed the recommendation of Anderson and Burnham (2002) and selected the most parsimonious model. Final models were validated by calculating variance inflation factors, inspecting residuals, testing for temporal and spatial autocorrelation, and evaluating model performance. We evaluated model goodness-of-fit following Nakagawa et al. (2017) for non-normal responses by calculating marginal and conditional delta-R^2^ using the *performance* package version 0.17.0 (Lüdecke et al. 2021). All statistical analyses were performed in R 4.5.2 (R Core Team, 2025).

## Results

### Context of Recorded Alarm Occurrence

Recorded alarm occurrence is a compound outcome that depends on encounter with a stimulus, its detection, the decision to call, and the event being observed and recorded. In a supplementary analysis (Appendix S1), the occurrence of recorded alarms to high-soaring species of bird varied strongly with season and daily maximum temperature, and more weakly with habitat, vegetation, and group size. These patterns are consistent with a large contribution of encounter and detection processes, although the analysis cannot separate these from the decision to call. We therefore did not use raw alarm occurrence as a measure of perceived risk, and conditioned our main analyses on a recorded alarm to a high-soaring bird.

### Do Meerkats Discriminate Between Predators and Non-Predators to Emit Alarm Calls?

For both seasonal threat discrimination analyses, two candidate models showed similar support (ΔAIC_c_ < 2; Tables S1.17 and S1.18). Therefore, we present the most parsimonious models’ results herein (Table 1), while the alternative models’ tables and plots are reported in supplementary material (Table S1.24; Figure S1.8). The proportion of alarm calls directed at predators was lower during the dry season (Figure 3), although this seasonal difference cannot be separated from seasonal changes in the relative abundance of vultures and eagles. Habitat differences in this response were smaller during the dry season (Figure 3A). As such, meerkats showed a lower probability to emit an alarm call in response to predators rather than non-predators during the wet season in high duneveld with 18.5% (95% CI: 14.7 – 22.9%) relative to the reference low duneveld habitat (24.1%; 21.1 – 27.3%; Figure 3A). In contrast, this probability weakly increased in low shrublands (28.3%; 24.4 – 32.7%) and riverbed and pans (27.0%; 23.7 – 30.6%; Figure 3A). During the dry season however, the threat discrimination probability only considerably differed from low duneveld (17.1%; 14.6 – 19.9%) in the riverbed and pans (20.8%; 17.7 – 24.1%), with a marginal increase in low shrublands (18.7; 15.4 – 22.5%; Figure 3A). Vegetation productivity showed contrasting habitat-specific effects. During the wet season, alarm calling to predators mainly decreased considerably in high duneveld dropping from 25.2% (19.7 – 31.8%) to 8.2% (3.6 – 17.4%) as NDVI increased from the 2.5^th^ to the 97.5^th^ percentiles of observed values (Figure 3C). We also observed an increase in low shrublands from 24.9% (20.5 – 29.9%) to 37.4% (28.6 – 47.1%; Figure 3E) and a weak marginal decrease in low duneveld from 25.4% (22.0 – 29.1%) to 21.1% (17.2 – 25.6%; Figure 3D). In contrast NDVI did not affect threat discrimination much in the riverbed and pans with a weak marginal decrease from 27.3% (23.8 – 31.2%) to 26.3% (22.0 – 31.0%; Figure 3B). During the dry season, low duneveld decreased from 18.7% (15.6 – 22.1%) to 14.6% (11.7 – 18.2%; Figure 3D), from which only high duneveld marginally differed with an increase from 14.1% (10.4 – 18.9%) to 21.2% (12.8 – 33.0%; Figure 3C). The within-season sensitivity analysis including ‘month’ (Appendix S1), however, indicated a less pronounced NDVI effect in low duneveld during both seasons. We detected effects of social contexts on threat discrimination, independently from habitat. During the wet season, the proportion of alarm calls directed at predators decreased from 25.6% (22.3 – 29.2%) for groups of 4 individuals to 21.5% (18.1 – 25.4%; Figure 3F) for groups of 28 individuals. However, the sensitivity analyses indicated this effect was not robust: it weakened when including ‘month’ changed sign when excluding the riverbed and pans (Appendix S1). During the dry season, however, the proportion of alarm calls directed at predators increased from 15.3% (12.8 – 18.0%) to 20.2% (16.8 – 24.1%; Figure 3F) with groups ranging from 4 to 32 individuals (2.5^th^ and 97.5^th^ percentiles). Furthermore, the presence of pups only marginally increased the proportion of alarm calls to predators from 24.1% (21.1 – 27.3%) to 24.9% (21.8 – 28.3%; Figure 3G). During the dry season, however, groups with pups directed 20.2% (17.1 – 23.4%) of alarm calls at predators, compared to 17.1% (14.6 – 19.9%) without pups, consistent with stronger threat discrimination (Figure 3G). Odds ratios are reported in Table S1.21.

**Figure 3.**
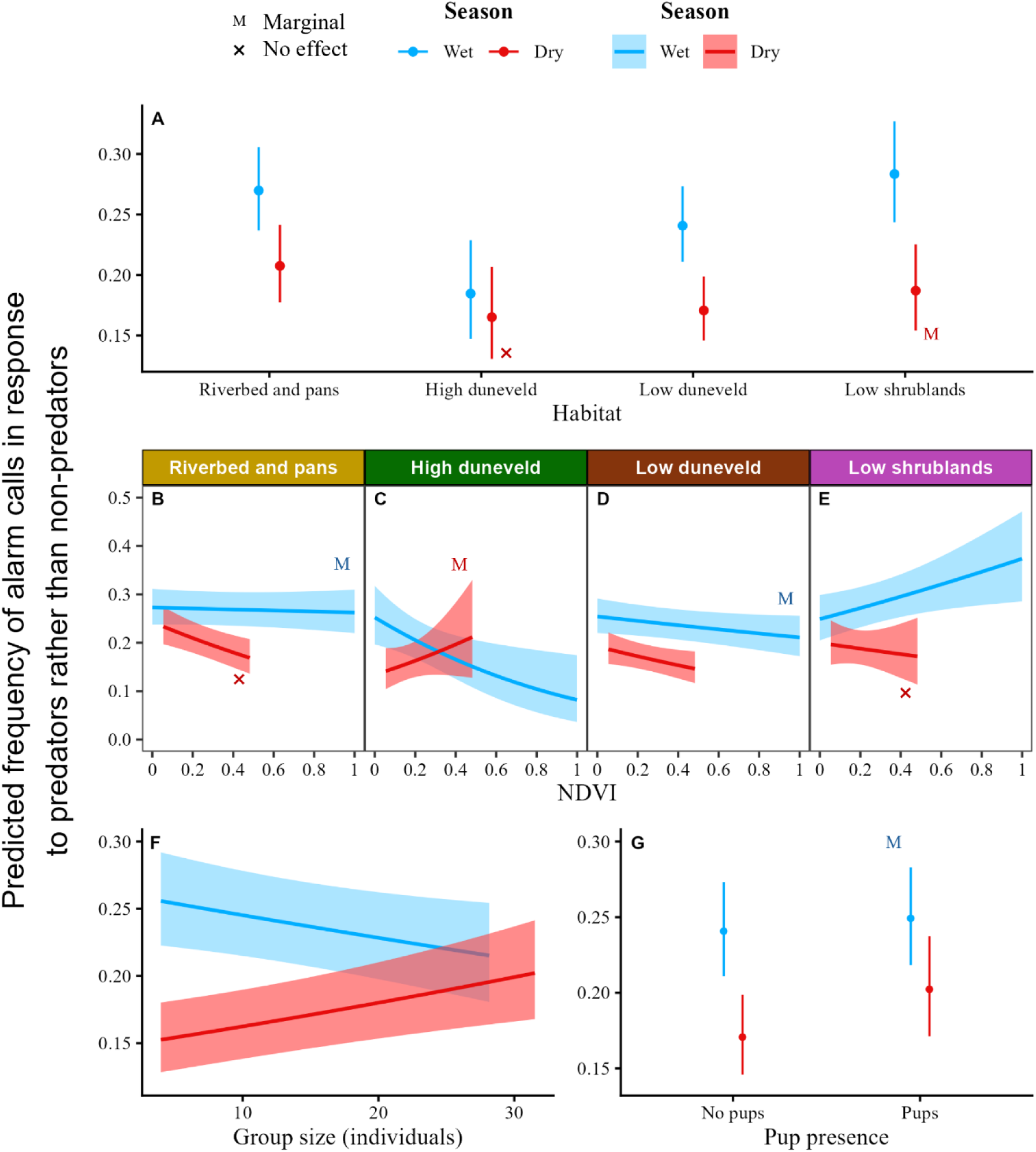
Predicted values for all the effects of the final model of the objective on threat discrimination for the wet (blue; marginal δ*R^2^*: 0.01; conditional δ*R^2^*: 0.05) and dry (red; marginal δ*R^2^*: 0.01; conditional δ*R^2^*: 0.07) seasons for which the 95% confidence intervals did not include 0. Error bars and ribbons are the 95% confidence intervals. To observe the effect better, y axes were scaled accordingly. (A) Baseline habitat structure. (B, C, D, E) Habitat-specific Normalised Difference Vegetation Index (NDVI) effects predicted over a more realistic NDVI range excluding 5% of extreme values, which was scaled to range between 0 and 1. (F) Group size effects predicted over a more realistic group size range excluding 5% of extreme values. (G) Pups.

**Table 1.** Estimates from the most parsimonious model for threat discrimination during both seasons. The wet and dry season models were fitted with 13,803 and 14,774 observations respectively, both from 54 meerkat groups over 23 years. Estimates where the confidence intervals excluded 0 are shown in bold. Abbreviations: SE, standard error; CI, confidence interval; σ², variance of random effects; NDVI, Normalised Difference Vegetation Index. Reference levels are low duneveld for habitat type, and no pups.

| Term | Wet season |  |  | Dry season |  |  |
| --- | --- | --- | --- | --- | --- | --- |
|  | Estimate | SE | 95% CI | Estimate | SE | 95% CI |
| **Fixed effects** |  |  |  |  |  |  |
| Intercept | **-1.149** | **0.087** | **-1.32 – -0.98** | **-1.581** | **0.095** | **-1.77 – -1.39** |
| *Spatial effects* |  |  |  |  |  |  |
| Riverbed and pans | **0.153** | **0.049** | **0.06 – 0.25** | **0.241** | **0.055** | **0.13 – 0.35** |
| High duneveld | **-0.337** | **0.116** | **-0.56 – -0.11** | -0.040 | 0.114 | -0.26 – 0.18 |
| Low shrublands | **0.221** | **0.073** | **0.08 – 0.36** | 0.111 | 0.085 | -0.06 – 0.28 |
| NDVI | -0.066 | 0.036 | -0.14 – 0 | **-0.077** | **0.039** | **-0.15 – 0** |
| Riverbed and pans × NDVI | 0.051 | 0.044 | -0.03 – 0.14 | -0.030 | 0.049 | -0.13 – 0.07 |
| High duneveld × NDVI | **-0.294** | **0.144** | **-0.58 – -0.01** | 0.205 | 0.108 | -0.01 – 0.42 |
| Low shrublands × NDVI | **0.224** | **0.074** | **0.08 – 0.37** | 0.034 | 0.087 | -0.14 – 0.2 |
| *Social effects* |  |  |  |  |  |  |
| Group size | **-0.056** | **0.026** | **-0.11 – -0.01** | **0.090** | **0.027** | **0.04 – 0.14** |
| Pups | 0.046 | 0.042 | -0.04 – 0.13 | **0.209** | **0.057** | **0.10 – 0.32** |
| **Random effects** |  |  |  |  |  |  |
| | $\sigma^2$ | | | $\sigma^2$ | | |
| Year | 0.110 |  |  | 0.126 |  |  |
| Group identity | 0.047 |  |  | 0.080 |  |  |

### Which Contexts Affect Response Escalation to Non-Predators?

The most supported model for the probability of collectively reacting with a strong rather than a weak response to a non-predator alarm call included spatial and social predictors, as well as an interaction between habitat type and pup presence (Table S1.19). Contrary to threat discrimination, the final model for reaction strength revealed no baseline habitat structure, with no significant difference from the 81.2% (74.2 – 86.7%) probability of the reference low duneveld habitat (Table 2; Figure 4A). There was, however, habitat-specific social context-dependence. Groups were more likely to display a strong reaction to an alarm call for a non-predator when foraging with pups in low duneveld with a 83.7% (77.2 – 88.6%) probability compared to 81.2% (74.2 – 86.7%) without pups (Figure 4C). Specifically, the probability of a strong reaction in riverbed and pans (81.9%; 75.0 – 87.3%) and low shrublands (81.1%; 73.7 – 86.9) increased for groups with pups to 87.3% (81.8 – 91.3%) and 87.8% (81.8 – 92.0%; Figure 4C). Moreover, increasing group size from 4 to 31 individuals (2.5^th^ to 97.5^th^ percentiles) raised the probability of responding with a strong reaction from 78.9% (71.2 – 85.0%) to 84.8% (78.5 – 89.5%; Figure 4B).

**Figure 4.**
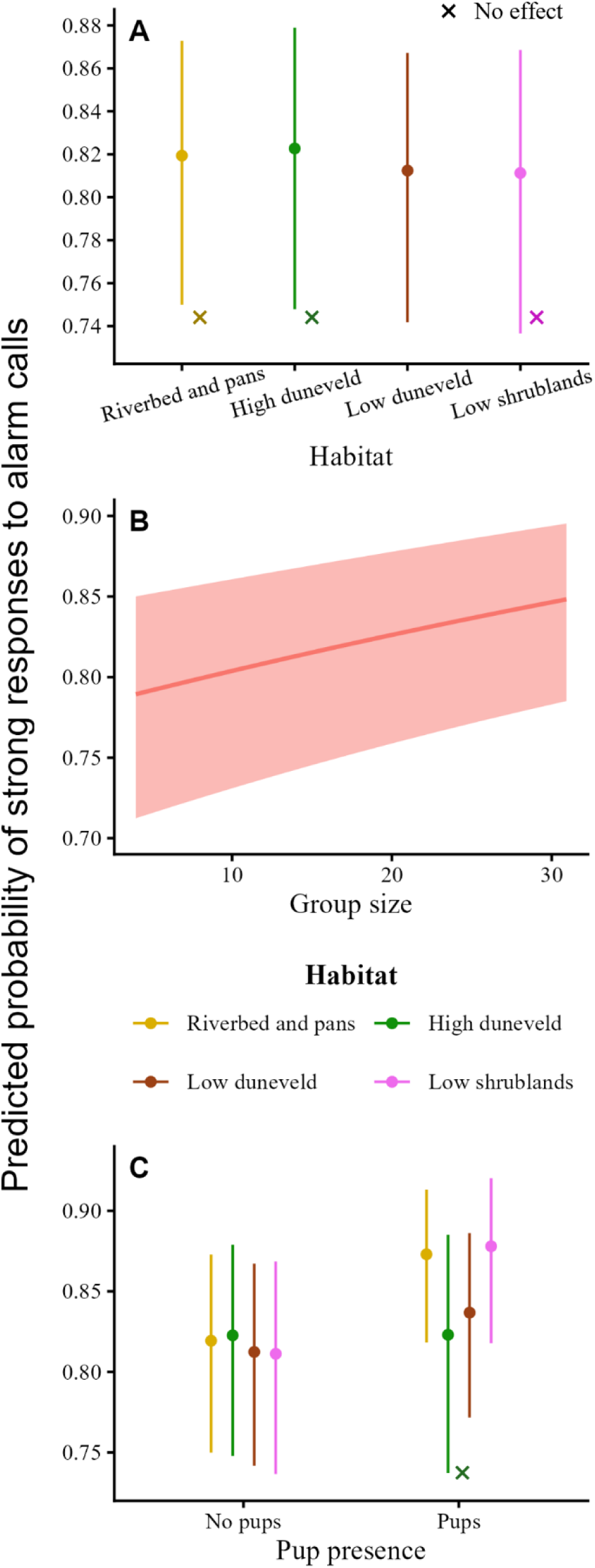
Predicted values for all the effects of the final model for response escalation with reaction strength for which the 95% confidence intervals did not include 0. Error bars and ribbons are the 95% confidence intervals. To observe the effect better, y axes were scaled accordingly. (A) Habitat baseline structure. (B) Group size effect predicted over a more realistic group size range excluding 5% of extreme values. (C) Habitat-specific effects of pups. Marginal δ*R^2^*: 0.08; conditional δ*R^2^*: 0.24.

**Table 2.** Estimates from the most parsimonious model for the probability of alarm calls eliciting strong reactions. The model was fitted with 21,598 observations from 55 groups over 23 years. Estimates where the confidence intervals excluded 0 are shown in bold. Abbreviations: SE, standard error; CI, confidence interval; σ², variance of random effects. Reference levels are low duneveld for habitat type, and no pups.

| Term | Estimate | SE | 95% CI |
| --- | --- | --- | --- |
| **Fixed effects** |  |  |  |
| Intercept | **1.466** | **0.209** | **1.06 – 1.88** |
| *Spatial effects* |  |  |  |
| Riverbed and pans | 0.046 | 0.052 | -0.05 – 0.15 |
| High duneveld | 0.069 | 0.102 | -0.13 – 0.27 |
| Low shrublands | -0.007 | 0.078 | -0.16 – 0.14 |
| *Social effects* |  |  |  |
| Group size | **0.100** | **0.021** | **0.06 – 0.14** |
| Pups | **0.169** | **0.055** | **0.06 – 0.28** |
| *Context-dependence* |  |  |  |
| Riverbed and pans × pups | **0.247** | **0.089** | **0.07 – 0.42** |
| High duneveld × pups | -0.167 | 0.178 | -0.52 – 0.18 |
| Low shrublands × pups | **0.346** | **0.151** | **0.05 – 0.64** |
| **Random effects** |  |  |  |
| | $\sigma^2$ | | |
| Year | 0.955 |  |  |
| Group identity | 0.043 |  |  |

For reaction intensity, two competing models could not be differentiated (Table S1.20). Thus, estimates of the alternative model can be found in supplementary (Table S1.25; Figure S1.9), while we present here the results of the most parsimonious model which only included habitat and social predictors (Table 3). Meerkats were more likely to flee relative to engaging in bipedal vigilance in reaction to a non-predator alarm call in the riverbed and pans (67.6%; 64.1 – 71.0%), high duneveld (67.4%; 62.8 – 71.7%), and low shrublands (63.8%; 59.5 – 68.0%; marginally), than groups in low duneveld (61.9%; 58.3 – 65.4%; Figure 5A). The probability of intense collective reactions also increased from 58.0% (54.0 – 62.0%) to 68.3% (64.1 – 72.1%) over group sizes ranging from 4 to 30 individuals (2.5^th^ to 97.5^th^ percentiles; Figure 5B). Odds ratios for reaction strength and intensity are reported in Tables S1.22 and S1.23.

**Figure 5.**
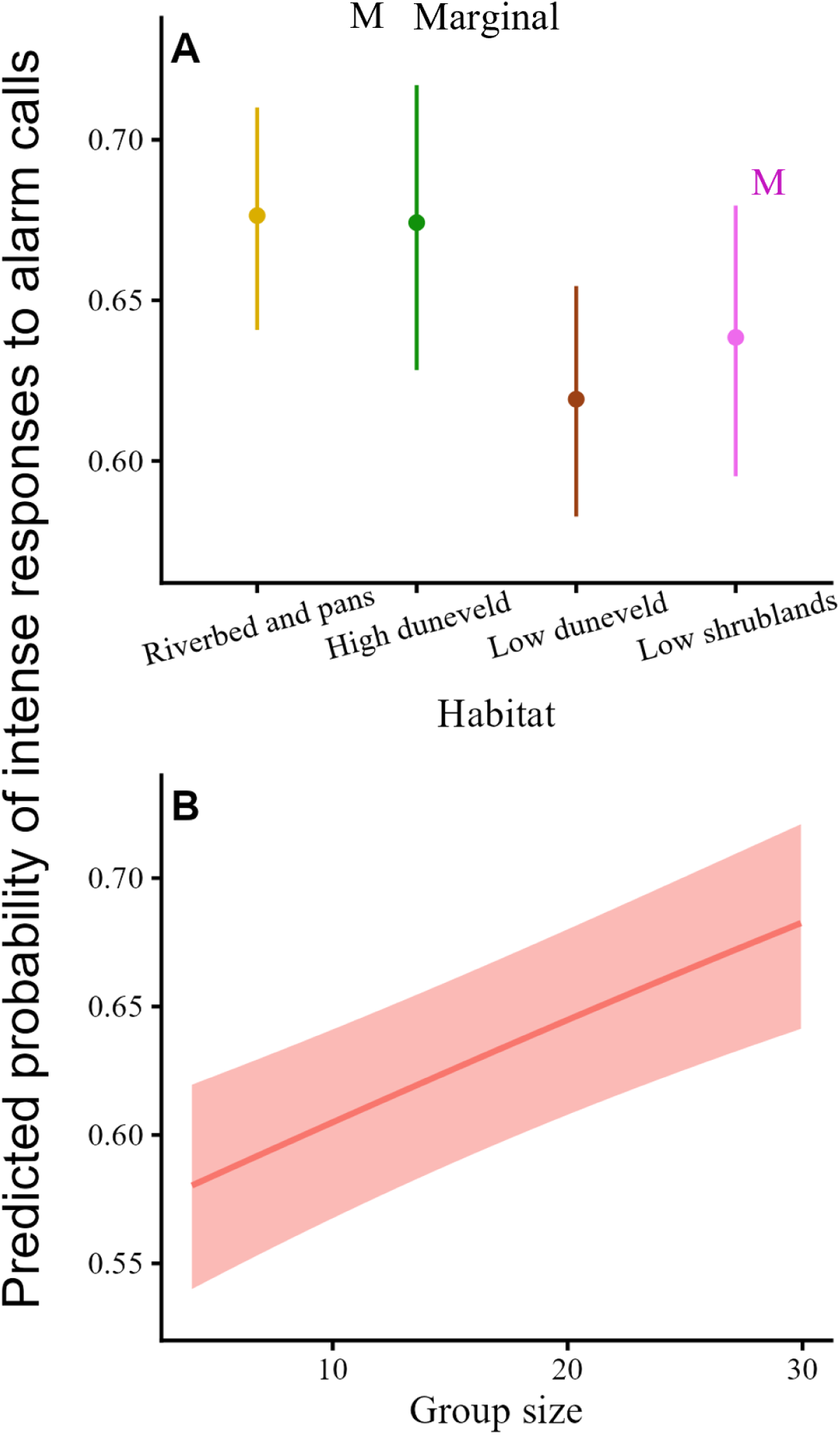
Predicted values for all the effects of the final model for response escalation with reaction intensity for which the 95% confidence intervals did not include 0. Error bars and ribbons are the 95% confidence intervals. To observe the effect better, y axes were scaled accordingly. (A) Baseline habitat structure. (B) Group size effect predicted over a more realistic group size range excluding 5% of extreme values. Marginal δ*R^2^*: 0.01; conditional δ*R^2^*: 0.05.

**Table 3.** Estimates from the most parsimonious model for the probability of alarm calls eliciting intense reactions (fleeing vs bipedal vigilance). The model was fitted with 17,063 observations from 55 groups over 23 years. Estimates where the confidence intervals excluded 0 are shown in bold. Abbreviations: SE, standard error; CI, confidence interval; σ², variance of random effects. Reference levels are low duneveld for habitat type, and no pups.

| Term | Estimate | SE | 95% CI |
| --- | --- | --- | --- |
| **Fixed effects** |  |  |  |
| Intercept | **0.486** | **0.078** | **0.33 – 0.64** |
| *Spatial effects* |  |  |  |
| Riverbed and pans | **0.251** | **0.041** | **0.17 – 0.33** |
| High duneveld | **0.241** | **0.076** | **0.09 – 0.39** |
| Low shrublands | 0.082 | 0.061 | -0.04 – 0.20 |
| *Social effects* |  |  |  |
| Group size | **0.110** | **0.021** | **0.07 – 0.15** |
| Pups | -0.024 | 0.035 | -0.09 – 0.04 |
| **Random effects** |  |  |  |
| | $\sigma^2$ | | |
| Year | 0.107 |  |  |
| Group identity | 0.024 |  |  |

## Discussion

In this study, we investigated the context-dependence of sequential sub-components of alarm calling in wild meerkats using 23 years of high-resolution behavioural data. Because alarm-calling-propensity is a compound outcome, we conditioned our main analyses on recorded alarm calls to high-soaring birds and examined predator–non-predator discrimination and the escalation of the ensuing collective responses to non-predators (Figure 1). The proportion of alarms directed at non-predators varied with some spatial and social contexts, and response escalation was associated mainly with social context, through group size and the presence of pups, although most effects were small. These findings highlight the relevance of decomposing behavioural response to risk rather than considering alarm calling as a simple behavioural process. Recorded alarm occurrence itself varied mainly with season and temperature (Appendix S1), consistent with a large contribution of encounter and detection processes. Conditioning on a recorded alarm reduces nuisance variation in total encounter, detection, and observation processes insofar as these affect eagle and vulture alarms similarly, but it does not remove species-specific differences in their availability or detectability; the predator–non-predator results should therefore be read as conditional on the assumption that such differences do not explain the context effects we report.

### Predator–Non-Predator Discrimination

Conditional on a recorded alarm, the proportion of alarms directed at non-predators varied with habitat, vegetation productivity, group size, and pup presence, but these effects were mostly small. Interpreting them as differences in discrimination rests on the assumption that the relative availability and detectability of eagles and vultures do not vary with the same predictors. This is least secure for habitat: the riverbed and pans include the fossil riverbed, which contains proportionally more large trees than the other habitats where both vultures and eagles perch and nest, so habitat effects may partly reflect where the birds are. Refugia density, although descriptive, may affect risk perception, which could contribute to the threat discrimination decrease in high duneveld and increase in low shrublands with more vegetation during the wet season.

### Social Context in Discrimination and Response Escalation

Group size had a small effect on discrimination during the dry season. Larger groups directed proportionally more alarms at predators, countering the largely accepted principle that large groups are expected to emit more false alarms (Beauchamp 2010; Gray and Webster 2023). While most studies focus on the advantages of larger groups (Prokopenko et al. 2024), this result may reflect costs of alarm calling associated with conspicuousness.

Pup presence influenced discrimination, albeit weakly. Although groups with pups have been shown to increase time allocation to vigilance (Santema and Clutton-Brock 2013), our dry season result indicate they instead directed proportionally fewer alarms at non-predators. As group with pups move more slowly (Thorley et al. 2026) and that pups do not react quickly to alarm calls, requiring protection (Clutton-Brock et al. 1999a), their presence increases collective vulnerability. Therefore, meerkats may respond to the higher bird abundances of dry months by prioritizing alarming at predators to avoid conspicuous reactions in the presence of pups. In this regard, Haff and Magrath (2013) showed that white-browed scrubwren (*Sericornis frontalis*) assess the vulnerability of their offspring to decide whether to alarm when predators are near their nest. Thus, threat discrimination could reflect social context-dependent decision-making in meerkats, although we emphasise that the pup effect was small.

Following non-predator alarms, response escalation was associated mainly with social context, particularly group size. Because this analysis excludes predators, whose alarms almost always elicited strong responses, it captures contextual variation in response thresholds towards a non-threatening stimulus, which is one component of perceived risk. The moderate effects of group size, particularly on reaction strength, are consistent with the stress transmission process via the social contagion of behavioural states (Engert et al. 2019; Brandl et al. 2022). As such, the percolation efficiency of wariness increases throughout the group with the number of individuals, facilitating the propagation of collective reactions (Brandl et al. 2022). Although not on the same collective reactions we considered in this paper, such social facilitation mechanism was shown experimentally in our population with meerkat displaying a considerably weaker mobbing response to playbacks directed at a few individuals, rather than conventional ones (Johnson-Ulrich et al. 2023). Thus, behavioural escalation may rely more on social contagion than on the acoustic properties of the signal itself, with larger groups being more prone to amplification cascades, in the absence of an actual predation threat (Tump et al. 2022). We do acknowledge, however, that the 50% recording threshold rule could contribute to the group size effects, because groups of different size may reach the threshold at different rates for weak and strong reactions.

Furthermore, groups with pups exhibited stronger reactions in the riverbed and pans and low shrublands, the habitats with the highest refugia densities (Appendix S1). In addition to raising conspicuousness, strong reactions, especially flight, can increase energy expenditure (Hall et al. 2013), but these costs may be alleviated when refugia can be reached rapidly. This is even more relevant for pups, for which being left behind by the group when tired is a leading cause of mortality (Russell et al. 2002). This pattern is consistent with a trade-off between social vulnerability and proximity to safety, although refugia densities remain descriptive, as we did not test it directly.

### Conclusions

In this study, we decomposed meerkat alarm calling behaviour in sequential components to evaluate how spatial, weather, and social contexts relate to predator–non-predator discrimination and response escalation, conditional on a recorded alarm. Together, our findings suggest that meerkat anti-predator behaviour may be context-dependent, although the effects we found were generally small. Social context, particularly group size, was associated with both components.

In a broader context, our results illustrate that a single observable anti-predator behaviour, such as alarm occurrence, need not reflect perceived risk directly. Our findings rather suggest that responses to the same risk can vary, depending on encounter probability, detection capacity, and perceived risk. Therefore, we recommend future risk perception studies to follow a similar approach by decomposing behavioural responses into sequential components, where possible, to avoid inferring risk perception from compound processes. Conditioning on observed events is useful, but each step carries its own assumptions, which should be stated and, where possible, tested.

Nevertheless, data collection for both predator and prey in natural systems is seldom feasible, leading studies on non-consumptive effects to use less reliable proxies (Moll et al. 2017; West et al. 2018; Prugh et al. 2019; Driscoll et al. 2022). Our study faced such limitation, as we lacked accurate estimates of predator abundance or occurrence. In particular, interpreting the predator–non-predator comparison as discrimination assumes that variation in the relative availability of vultures and eagles does not explain the context effects we report, which independent raptor surveys could test. Rather than assuming consistent encounter probability, we instead separated the analyses seasonally to account for breeding periods. Thereby, our work presents a case study using a process-based approach which complements existing proxies to improve inference on prey responses to risk.

## Acknowledgments

We thank the Kalahari Research Trust and the Kalahari Meerkat project for hosting this study, long-term data collection and habituation of the meerkats. We are further thankful for the landowners of the properties neighbouring the Kalahari Research Centre for allowing the monitoring of meerkats onto their lands. We express a deep gratitude to all the volunteers, research assistants, students, managers, researchers, and staff members who made this project possible. Particularly, we thank Walter Jubber, David Gaynor and Tim Vink for management of the reserve and the project’s databases. We further thank the Mammal Research Institute and the animal ethics committee at the University of Pretoria for their logistical support, as well as the Northern Cape Department of Environment and Nature Conservation for granting permission for this research. The long-term data collection was permitted by the University of Pretoria’s animal ethics committee (NAS003/2022, EC047-16, EC010-13, SOP029-12) and the Northern Cape Conservation Service, South Africa (FAUNA 0930/2022, FAUNA 1020/2016, FAUNA 192/2014, FAUNA 050/2013, FAUNA 263/3/2012) and followed the Treatment of Animals in Behavioural Research and Teaching standards outlined by ASAB/ABS (2012).

## Conflict of interest

The authors declare no conflicts of interest.

## Author contributions

Zachary Cloutier: Conceptualization, Methodology, Formal analysis, Data curation, Visualization, Writing – original draft, Writing – review & editing. Marta B. Manser: Conceptualization, Methodology, Data curation, Funding acquisition, Project administration, Supervision, Writing – review & editing. Jack Thorley: Writing – review & editing. Dominik M. Behr: Writing – review & editing. Tim H. Clutton-Brock: Data curation, Funding acquisition, Project administration, Writing – review & editing. Arpat Ozgul: Conceptualization, Methodology, Funding acquisition, Supervision, Writing – review & editing.

## Funding

The long-term meerkat research project is funded by the European Union’s Horizon 2020 research and innovation program from the European Research Council (no. 742808 and no. 294494 to T.H. Clutton-Brock), the Human Frontier Science Program (no. RGP0051/2017 and no. RGP0051/2019), the University of Cambridge (through T.H. Clutton-Brock), the University of Zurich (through M.B. Manser), the Newton Trust, the Mava Foundation, the Zoo Zurich, the Exekias foundation, and the Irene Staehelin foundation. Z. Cloutier’s salary and field costs were funded through UZH faculty funding provided to A. Ozgul.

## Notes

### Competing Interest Statement

The authors have declared no competing interest.

