## Supplementary Material: Appendix S1 for "Decomposing alarm-calling behaviour to investigate context-dependent threat discrimination and response escalation in a social mongoose"

### Sections

#### 1. Meerkat Alarm Calls

**Table S1.1.** Classifications of the stimuli documented to have triggered alarm calls in meerkats between 2002 and 2024. The observations in the "Unknown" category did not carry enough information to classify them as a threat or not to meerkats. Representative taxa are listed for each species type. The count and percentage of the total alarm call count is listed for each species and each category.

| Category | Type | Representative taxa | Count | % |
|---|---|---|---|---|
| Threats 10,660 (4.5%) | Soaring aerial | Eagles | 6,633 | 2.80 |
| | Non-soaring aerial | Falcons & owls | 3,311 | 1.40 |
| | Snakes | Cobras & adders | 579 | 0.25 |
| | Mammals | Jackals, felines & hyenas | 137 | 0.06 |
| Non-threats 116,474 (49.3%) | Soaring aerial | Vultures | 22,636 | 9.59 |
| | Non-soaring aerial | Kites & kestrels | 15,320 | 6.48 |
| | Insectivores | Foxes & mongooses | 1,625 | 0.69 |
| | Herbivores | Ungulates & hares | 9,550 | 4.04 |
| | Extra-specific alarms | Birds & ground squirrels | 44,680 | 18.92 |
| | Secondary cues | Objects & scents | 3,730 | 1.58 |
| | Anthropogenic stimuli | Humans & vehicles | 17,246 | 7.30 |
| | False alarms | Wind & noises | 1,687 | 0.71 |
| Unknown 109 024 (46.2%) | Aerial alarms | | 23,314 | 9.87 |
| | Terrestrial alarms | | 636 | 0.27 |
| | Unknown | | 85,074 | 36.02 |
| | | **Total** | **236,158** | |

**Table S1.2.** Lists of all alarm calls and their stimuli between 2002 and 2024 classified as known threats or non-threats for meerkats.

| Trigger | | | Strong | | | Weak |
|---|---|---|---|---|---|---|
| **Category** | **Type** | **Species** | **Fleeing (Intense)** | **Vigilance (Moderate)** | **Mobbing** | **Look briefly** |
| Threats 10660 (4.5%) | Soaring birds 6633 (2.8%) | Martial eagle 3791 (1.6%) | 2908 (76.7%) | 763 (20.1%) | 4 (0.1%) | 66 (1.7%) |
| | | Tawny eagle 456 (0.2%) | 323 (70.8%) | 90 (19.7%) | 1 (0.2%) | 25 (5.5%) |
| | | Steppe eagle 34 (0%) | 16 (47.1%) | 13 (38.2%) | 1 (2.9%) | 4 (11.8%) |
| | | Wahlberg's eagle 33 (0%) | 26 (78.8%) | 3 (9.1%) | 0 (0%) | 4 (12.1%) |
| | | Booted eagle 13 (0%) | 9 (69.2%) | 3 (23.1%) | 0 (0%) | 0 (0%) |
| | | Und. eagle 108 (0%) | 73 (67.6%) | 27 (25%) | 0 (0%) | 2 (1.9%) |
| | | Black-breasted snake eagle 1804 (0.8%) | 1296 (71.8%) | 431 (23.9%) | 3 (0.2%) | 47 (2.6%) |
| | | Brown snake eagle 84 (0%) | 58 (69%) | 22 (26.2%) | 1 (1.2%) | 1 (1.2%) |
| | | Bateleur 310 (0.1%) | 207 (66.8%) | 59 (19%) | 4 (1.3%) | 39 (12.6%) |
| | Non-soaring birds 3311 (1.4%) | Pale chanting goshawk 2482 (1.1%) | 1532 (61.7%) | 695 (28%) | 47 (1.9%) | 124 (5%) |
| | | Lanner falcon 153 (0.1%) | 77 (50.3%) | 62 (40.5%) | 3 (2%) | 9 (5.9%) |
| | | Peregrine falcon 14 (0%) | 10 (71.4%) | 2 (14.3%) | 2 (14.3%) | 0 (0%) |
| | | Red-necked falcon 8 (0%) | 4 (50%) | 3 (37.5%) | 0 (0%) | 1 (12.5%) |
| | | Und. falcon/kestrel 230 (0.1%) | 145 (63%) | 62 (27%) | 0 (0%) | 16 (7%) |
| | | Owl 63 (0%) | 39 (61.9%) | 17 (27%) | 7 (11.1%) | 0 (0%) |
| | | Spotted eagle owl 37 (0%) | 10 (27%) | 5 (13.5%) | 6 (16.2%) | 0 (0%) |
| | | Barn owl 39 (0%) | 26 (66.7%) | 9 (23.1%) | 4 (10.3%) | 0 (0%) |
| | | Giant eagle owl 79 (0%) | 39 (49.4%) | 32 (40.5%) | 2 (2.5%) | 4 (5.1%) |
| | | Secretary bird 25 (0%) | 20 (80%) | 4 (16%) | 0 (0%) | 1 (4%) |
| | | Montagu's harrier 9 (0%) | 5 (55.6%) | 2 (22.2%) | 2 (22.2%) | 0 (0%) |
| | | African marsh harrier 1 (0%) | 1 (100%) | 0 (0%) | 0 (0%) | 0 (0%) |
| | | Gymnogene 1 (0%) | 0 (0%) | 1 (100%) | 0 (0%) | 0 (0%) |
| | | Little banded goshawk 5 (0%) | 2 (40%) | 1 (20%) | 0 (0%) | 0 (0%) |
| | | Steppe buzzard 165 (0.1%) | 98 (59.4%) | 45 (27.3%) | 3 (1.8%) | 17 (10.3%) |
| | Snakes 579 (0.2%) | Snake 150 (0.1%) | 3 (2%) | 8 (5.3%) | 132 (88%) | 0 (0%) |
| | | Cape cobra 103 (0%) | 7 (6.8%) | 1 (1%) | 89 (86.4%) | 1 (1%) |
| | | Puff adder 251 (0.1%) | 4 (1.6%) | 1 (0.4%) | 229 (91.2%) | 2 (0.8%) |
| | | Horned adder 75 (0%) | 0 (0%) | 2 (2.7%) | 58 (77.3%) | 0 (0%) |
| | Mammals 137 (0.1%) | Black-backed jackal 14 (0%) | 9 (64.3%) | 4 (28.6%) | 1 (7.1%) | 0 (0%) |
| | | African wild cat 40 (0%) | 18 (45%) | 7 (17.5%) | 15 (37.5%) | 0 (0%) |
| | | Small spotted cat 8 (0%) | 2 (25%) | 0 (0%) | 6 (75%) | 0 (0%) |
| | | Striped polecat 23 (0%) | 3 (13%) | 0 (0%) | 20 (87%) | 0 (0%) |
| | | Domestic cat 21 (0%) | 7 (33.3%) | 8 (38.1%) | 5 (23.8%) | 1 (4.8%) |
| | | Feline 3 (0%) | 1 (33.3.%) | 0 (0%) | 2 (66.7%) | 0 (0%) |
| | | Brown hyena 3 (0%) | 1 (33.3%) | 2 (66.7%) | 0 (0%) | 0 (0%) |
| | | Small spotted genet 24 (0%) | 1 (4.2%) | 1 (4.2%) | 22 (91.7%) | 0 (0%) |
| | | Baboon 1 (0%) | 1 (100%) | 0 (0%) | 0 (0%) | 0 (0%) |
| Non-threats 116474 (49.3%) | Soaring birds 22636 (9.6%) | White-backed vulture 17769 (7.5%) | 8483 (47.7%) | 5116 (28.8%) | 38 (0.2%) | 3762 (21.2%) |
| | | Lappet-faced vulture 3589 (1.5%) | 1978 (55.1%) | 1002 (27.9%) | 2 (0.1%) | 540 (15%) |
| | | Cape vulture 107 (0%) | 50 (46.7%) | 19 (17.8%) | 1 (0.9%) | 26 (24.3%) |
| | | White-headed vulture 23 (0%) | 15 (65.2%) | 7 (30.4%) | 0 (0%) | 1 (4.3%) |
| | | Und. vulture 1148 (0.5%) | 521 (45.3%) | 282 (24.6%) | 0 (0%) | 266 (23.2%) |
| | Non-soaring birds 15320 (6.5%) | Non-threatening bird 13210 (5.6%) | 7203 (54.5%) | 3012 (22.8%) | 96 (0.7%) | 2554 (19.3%) |
| | | Guinea fowl 78 (0%) | 43 (55.1%) | 19 (24.4%) | 1 (1.3%) | 13 (16.7%) |
| | | Black kite 166 (0.1%) | 117 (70.5%) | 44 (26.5%) | 0 (0%) | 5 (3%) |
| | | Yellow-billed kite 4 (0%) | 0 (0%) | 2 (50%) | 0 (0%) | 1 (25%) |
| | | Kori bustard 244 (0.1%) | 171 (70.1%) | 38 (15.6%) | 1 (0.4%) | 24 (9.8%) |
| | | Pied babbler 1 (0%) | 1 (100%) | 0 (0%) | 0 (0%) | 0 (0%) |
| | | Black-shouldered kite 752 (0.3%) | 457 (60.8%) | 239 (31.8%) | 4 (0.5%) | 41 (5.5%) |
| | | Gabar goshawk 243 (0.1%) | 136 (56%) | 66 (27.2%) | 2 (0.8%) | 32 (13.2%) |
| | | Pygmy falcon 60 (0%) | 38 (63.3%) | 10 (16.7%) | 0 (0%) | 11 (18.3%) |
| | | Pearl spotted owl 6 (0%) | 5 (83.3%) | 0 (0%) | 0 (0%) | 1 (16.7%) |

| Trigger | | | Strong | | | Weak |
|---|---|---|---|---|---|---|
| **Category** | **Type** | **Species** | **Fleeing (Intense)** | **Vigilance (Moderate)** | **Mobbing** | **Look briefly** |
| | | Hobby falcon 3 (0%) | 2 (66.7%) | 1 (33.3%) | 0 (0%) | 0 (0%) |
| | | Rock kestrel 13 (0%) | 13 (100%) | 0 (0%) | 0 (0%) | 0 (0%) |
| | | Black harrier 2 (0%) | 1 (50%) | 1 (50%) | 0 (0%) | 0 (0%) |
| | | Scops owl 1 (0%) | 0 (0%) | 0 (0%) | 0 (0%) | 1 (100%) |
| | | Small und. raptor 500 (0.2%) | 289 (57.8%) | 139 (27.8%) | 4 (0.8%) | 65 (13%) |
| | | Greater kestrel 25 (0%) | 14 (56%) | 8 (32%) | 0 (0%) | 3 (12%) |
| | | Lesser kestrel 11 (0%) | 11 (100%) | 0 (0%) | 0 (0%) | 0 (0%) |
| | | Bat 1 (0%) | 0 (0%) | 0 (0%) | 1 (100%) | 0 (0%) |
| | Insectivores 1625 (0.7%) | Meerkat 25 (0%) | 13 (52%) | 9 (36%) | 1 (4%) | 2 (8%) |
| | | Yellow mongoose 320 (0.1%) | 51 (15.9%) | 105 (32.8%) | 123 (38.4%) | 27 (8.4%) |
| | | Slender mongoose 185 (0.1%) | 28 (15.1%) | 88 (47.6%) | 32 (17.3%) | 27 (14.6%) |
| | | Mongoose 2 (0%) | 0 (0%) | 2 (100%) | 0 (0%) | 0 (0%) |
| | | Cape fox 121 (0.1%) | 29 (24%) | 42 (34.7%) | 47 (38.8%) | 2 (1.7%) |
| | | Bat-eared fox 447 (0.2%) | 188 (42.1%) | 183 (40.9%) | 62 (13.9%) | 11 (2.5%) |
| | | Fox 1 (0%) | 0 (0%) | 1 (100%) | 0 (0%) | 0 (0%) |
| | | Aardvark 33 (0%) | 18 (54.5%) | 7 (21.2%) | 3 (9.1%) | 5 (15.2%) |
| | | Aardwolf 21 (0%) | 14 (66.7%) | 4 (19%) | 2 (9.5%) | 1 (4.8%) |
| | | Warthog 12 (0%) | 8 (66.7%) | 4 (33.3%) | 0 (0%) | 0 (0%) |
| | | Lizard 117 (0%) | 2 (1.7%) | 12 (10.3%) | 101 (86.3%) | 0 (0%) |
| | | Ostrich 340 (0.1%) | 178 (52.4%) | 105 (30.9%) | 3 (0.9%) | 52 (15.3%) |
| | | Pangolin 1 (0%) | 0 (0%) | 1 (100%) | 0 (0%) | 0 (0%) |
| | Herbivores 9550 (4%) | Tortoise 88 (0%) | 5 (5.7%) | 21 (23.9%) | 54 (61.4%) | 4 (4.5%) |
| | | Steenbuck 971 (0.4%) | 328 (33.8%) | 390 (40.2%) | 27 (2.8%) | 188 (19.4%) |
| | | Springbok 1331 (0.6%) | 386 (29%) | 598 (44.9%) | 21 (1.6%) | 271 (20.4%) |
| | | Hare (464 (0.2%) | 83 (17.9%) | 133 (28.7%) | 165 (35.6%) | 60 (12.9%) |
| | | Grey duiker 489 (0.2%) | 169 (34.6%) | 198 (40.5%) | 6 (1.2%) | 96 (19.6%) |
| | | Blue wildebeest 252 (0.1%) | 81 (32.1%) | 125 (49.6%) | 1 (0.4%) | 37 (14.7%) |
| | | Eland 330 (0.1%) | 122 (37%) | 137 (41.5%) | 4 (1.2%) | 54 (16.4%) |
| | | Gemsbok 99 (0%) | 22 (22.2%) | 55 (55.6%) | 2 (2%) | 16 (16.2%) |
| | | Red hartebeest 100 (0%) | 38 (38%) | 46 (46%) | 0 (0%) | 13 (13%) |
| | | Waterbuck 12 (0%) | 3 (25%) | 9 (75%) | 0 (0%) | 0 (0%) |
| | | Roan antelope 3 (0%) | 0 (0%) | 3 (100%) | 0 (0%) | 0 (0%) |
| | | Giraffe 2 (0%) | 1 (50%) | 1 (50%) | 0 (0%) | 0 (0%) |
| | | Horse/rider 2041 (0.9%) | 981 (48.1%) | 690 (33.8%) | 10 (0.5%) | 293 (14.4%) |
| | | Camel 1 (0%) | 1 (100%) | 0 (0%) | 0 (0%) | 0 (0%) |
| | | Antelope 15 (0%) | 2 (13.3%) | 8 (53.3%) | 1 (6.7%) | 3 (20%) |
| | | Porcupine 3 (0%) | 0 (0%) | 2 (66.7%) | 1 (33.3%) | 0 (0%) |
| | | Stock 3245 (1.4%) | 1370 (42.2%) | 1217 (37.5%) | 34 (1%) | 543 (16.7%) |
| | | Small mammal 88 (0%) | 31 (35.2%) | 38 (43.2%) | 11 (12.5%) | 8 (9.1%) |
| | | Ground squirrel 16 (0%) | 6 (37.5%) | 4 (25%) | 4 (25%) | 2 (12.5%) |
| | Extra-specific alarm calls 44680 (18.9%) | Bird 35216 (14.9%) | 21834 (62%) | 6406 (18.2%) | 71 (0.2%) | 5936 (16.9%) |
| | | Drongo 9250 (3.9%) | 6918 (74.8%) | 1034 (11.2%) | 24 (0.3%) | 1189 (12.9%) |
| | | Ground squirrel 211 (0.1%) | 69 (32.7%) | 54 (25.6%) | 35 (16.6%) | 39 (18.5%) |
| | | Other meerkat 3 (0%) | 1 (33.3%) | 2 (66.7%) | 0 (0%) | 0 (0%) |
| | Secondary cues 3730 (1.6%) | Scent 3178 (1.3%) | 19 (0.6%) | 9 (0.3%) | 3042 (95.7%) | 3 (0.1%) |
| | | Fur/faeces/feathers 547 (0.2%) | 81 (14.8%) | 18 (3.3%) | 405 (74%) | 15 (2.7%) |
| | | Carcass 5 (0%) | 0 (0%) | 0 (0%) | 5 (100%) | 0 (0%) |
| | Anthropogenic stimuli 17246 (7.3%) | Human 4196 (1.8%) | 2052 (48.9%) | 1359 (32.4%) | 14 (0.3%) | 651 (15.5%) |
| | | Gun shot 40 (0%) | 23 (57.5%) | 11 (27.5%) | 0 (0%) | 6 (15%) |
| | | Dog 1800 (0.8%) | 976 (54.2%) | 608 (33.8%) | 9 (0.5%) | 187 (10.4%) |
| | | Horse-drawn vehicle 221 (0.1%) | 124 (56.1%) | 59 (26.7%) | 0 (0%) | 32 (14.5%) |
| | | Motor vehicle 10546 (4.5%) | 5551 (52.6%) | 2849 (27%) | 7 (0.1%) | 1876 (17.8%) |
| | | Drone 1 (0%) | 1 (100%) | 0 (0%) | 0 (0%) | 0 (0%) |
| | | Helicopter 19 (0%) | 5 (26.3%) | 8 (42.1%) | 0 (0%) | 6 (31.6%) |
| | | Airplane 423 (0.2%) | 270 (63.8%) | 105 (24.8%) | 2 (0.5%) | 41 (9.7%) |
| | False alarms 1687 (0.7%) | Thunder 149 (0.1%) | 100 (67.1%) | 24 (16.1%) | 1 (0.7%) | 22 (14.8%) |
| | | Noise 72 (0%) | 44 (61.1%) | 19 (26.4%) | 0 (0%) | 6 (8.3%) |

| Trigger | | | Strong | | | Weak |
|---|---|---|---|---|---|---|
| **Category** | **Type** | **Species** | **Fleeing (Intense)** | **Vigilance (Moderate)** | **Mobbing** | **Look briefly** |
| | | Wind 1466 (0.6%) | 1119 (76.3%) | 254 (17.3%) | 1 (0.1%) | 88 (6%) |
| Unknown 109024 (46.2%) | Und birds 23314 (9.9%) | Aerial alarm 164 (0.1%) | 95 (57.9%) | 27 (16.5%) | 0 (0%) | 42 (25.6%) |
| | | L und. raptor 10044 (4.3%) | 5974 (59.5%) | 2833 (28.2%) | 23 (0.2%) | 1055 (10.5%) |
| | | M und. raptor 1953 (0.8%) | 1141 (58.4%) | 582 (29.8%) | 2 (0.1%) | 203 (10.4%) |
| | | Und. raptor 11153 (4.7%) | 6802 (61%) | 2436 (21.8%) | 12 (0.1%) | 1515 (13.6%) |
| | Und terrestrial 636 (0.3%) | Terrestrial alarm 636 (0.3%) | 299 (47%) | 291 (45.8%) | 15 (2.4%) | 31 (4.9%) |
| | Unknowns 85074 (36%) | Unknown 85074 (36%) | 55820 (65.6%) | 15526 (18.2%) | 1019 (1.2%) | 10212 (12%) |
| Total 236158 (100%) | | - 236158 (100%) | 140005 (59.3%) | 51142 (21.7%) | 6221 (2.6%) | 32580 (13.8%) |

#### 2. Habitats

**Table S1.3.** Habitat classification of the Kalahari Research Centre, based on soil and vegetation characteristics.

| Habitat | Description |
|---|---|
| Riverbed and pans | Compound category of the dry fossil riverbed and terraces as well as calcareous pans and flats. This category is characterised by calcareous soil, open woodland and tree savannah with sparse grasses, shrubs, and forbs (van Rooyen et al. 2008). |
| High duneveld | Grass-dominated plains with sparse shrubs with *Stipagrostis amabilis* as indicator species (van Rooyen et al. 2008). |
| Low duneveld | Mixed acacia savannah on red sand where the spatial density of woody species, grasses, sedges, dwarf shrubs, and forbs varies (van Rooyen et al. 2008). |
| Low shrublands | Habitat defined by a moderate cover of dense drie doring (*Rhigozum trichotomum*). |

#### 3. Hypotheses and Predictions

**Table S1.4.** Predictions and mechanisms for each variable and interaction terms. Hyp is hypothesis.

| Hyp | Interaction | Prediction and mechanisms |
|---|---|---|
| **Spatial contexts** | Habitat | We predicted a lower *threat discrimination* and greater *response escalation* in habitats with lower refugia densities (Rosenzweig 1974; Camp et al. 2012). |
| | NDVI | *Threat discrimination* should increase as thicker vegetation would provide safety through concealment. |
| | Habitat x NDVI | Vegetation effect on *threat discrimination* should be lesser in the riverbed and pans with the lack of vegetation, providing no cover for meerkats. It should also be weaker in low shrublands as branches already provide structural protection. |
| **Weather contexts** | Visibility | *Threat discrimination* should decrease with impaired visual detection causing an increase in uncertainty. |
| | Acoustics | *Response escalation* should be greater with impaired information transmission, causing more uncertainty (Hayes and Huntly 2005; Beauchamp 2010). |
| **Social contexts** | Group size | Larger groups should *discriminate* better due to increased perceived safety associated with confusion and dilution effects (Krakauer 1995; Lehtonen and Jaatinen 2016). *Response escalation* should be greater in larger groups due to social contagion (Brandl et al. 2022) |
| | Pups | *Threat discrimination* should be greater and *collective reactions* stronger and more intense in groups with pups due to increased vulnerability (Lima and Dill 1990). |
| **Habitat-specific context-dependence** | x Visibility | Poor visibility conditions should decrease the relative habitat-specific differences in *threat discrimination* as open habitats should not confer a visual advantage over denser habitats. |
| | x Acoustics | The effect of poor acoustic conditions on *response escalation* should be more pronounced in low shrublands as visibility of both conspecifics and stimuli should already be obstructed. |
| | x Group size | The effect of group size on *threat discrimination* should be higher in the riverbed and pans, where the conspicuousness should be more pronounced. |
| | x Pups | The vulnerability of groups with pups should be heightened in open habitats, thereby lowering the *discrimination of aerial stimuli* and increasing the likelihood of displaying *strong and intense reactions*. |

#### 4. Habitat-NDVI Correlations

As NDVI and habitat may be correlated, we investigated this relationship between the two variables to have a clear idea of the situation. First, we looked at the mean NDVI for each habitat type within the buffer zone over the entire study period.

**Table S1.5.** NDVI descriptive statistics for all locations between 2002 and 2024. IQR is the interquartile range.

| Season | Habitat | NDVI | SD | Median | IQR | Locations |
|---|---|---|---|---|---|---|
| Dry | Low duneveld | 1,949.9 | 312.27 | 1,886 | 344 | 73,642 |
| | Riverbed and pans | 1,937.7 | 232.89 | 1,896 | 278 | 83,693 |
| | Low shrublands | 1,857.0 | 239.90 | 1,823 | 290 | 16,592 |
| | High duneveld | 1,832.5 | 217.43 | 1,794 | 273 | 10,785 |
| Wet | Low duneveld | 2,077.9 | 534.28 | 1,886 | 531 | 119,834 |
| | Riverbed and pans | 2,038.0 | 654.14 | 1,822 | 651 | 110,337 |
| | Low shrublands | 2,028.6 | 603.37 | 1,833 | 726 | 26,554 |
| | High duneveld | 1,890.6 | 450.43 | 1,714 | 446 | 14,560 |

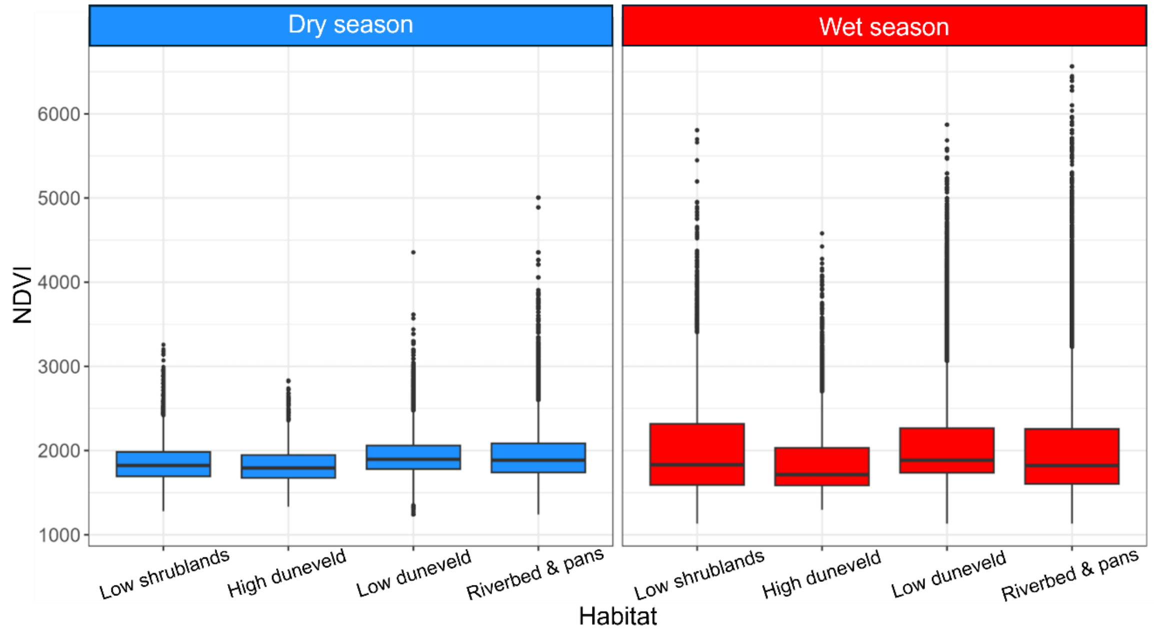

**Figure S1.1.** Boxplots of the distribution of NDVI values per habitat per season between 2002 and 2024.

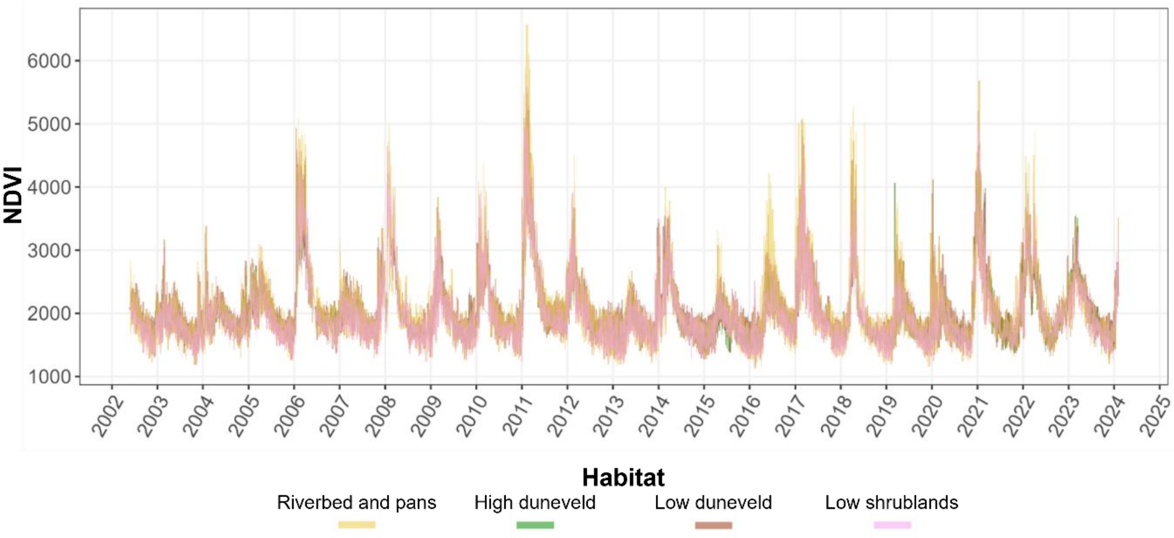

**Figure S1.2.** Time-series of all NDVI values during the temporal range of our study. The four lines are the NDVI values for each of the four habitats.

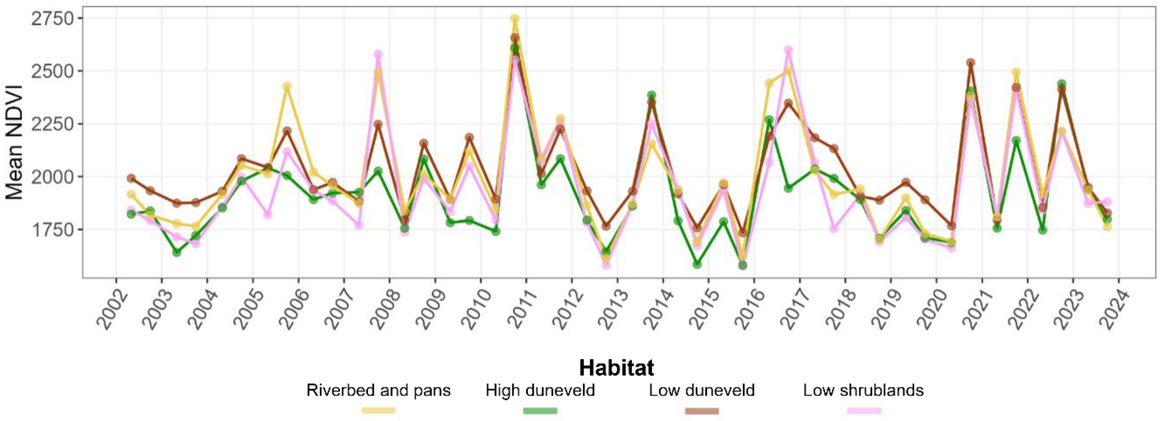

**Figure S1.3.** Time-series of seasonal NDVI mean values. The four lines are the NDVI values for each of the four habitats. Each dot is a season mean value.

**Table S1.6.** Results of the Kruskal-Wallis tests to differentiate NDVI values between habitats all at once and when separating by season.

| Season | Statistic | Effect size | Magnitude | P-value |
|---|---|---|---|---|
| Dry | 3,624.091 | 0.020 | small | <0.001 |
| Wet | 5,401.225 | 0.020 | small | <0.001 |

Note: Statistically there is a difference, but it is not meaningful.

We further converted habitat types into binary dummy variables to compute point-biserial correlations between NDVI and each habitat type. Point-biserial correlation is mathematically equivalent to Pearson correlation between one continuous and one binary variable.

**Table S1.7.** Point-biserial correlation between each habitat type and NDVI for all values at once as well as per season.

| Season | Habitat | Correlation |
|---|---|---|
| Dry | Riverbed and pans | 0.06 |
| | Low Duneveld | 0.03 |
| | Low shrublands | -0.08 |
| | High duneveld | **-0.09** |
| Wet | Low Duneveld | 0.05 |
| | Low shrublands | -0.01 |
| | Riverbed and pans | -0.01 |
| | High duneveld | **-0.06** |

#### 5. Refugia Analysis

##### *5A - Refugia Methods*

Because we did not have continuous documentation of boltholes for the entire span of the study, we determined whether habitats differed in their accessibility to refugia (boltholes and sleeping burrows). To achieve this, we quantified the spatial distribution of refugia across habitat types. First, we calculated 95% kernel density estimates (KDEs; Matthiopoulos et al. 2020; Parsons et al. 2022; Gil et al. 2025) for each group-season-year combination with at least 20 observations using the *amt* package version 0.3.1.0 (Signer et al. 2019).We then extracted the entire combined area used by meerkats between 2002 and 2024 for each habitat type and calculated refugia densities. To test whether refugia densities were proportionally distributed among habitats, we compared observed and expected bolthole, sleeping burrow, and all refugia densities separately using $\chi^2$ goodness-of-fit tests.

##### *5B - Refugia Results*

We quantified the spatial distribution of refugia to assess whether refugia accessibility differed among habitats (Table S1.8; Figure S1.4). We found that although low duneveld had the largest area and the most refugia (5,390; 45.5%), the riverbed and pans was the habitat with the highest refuge density (218.4 refugia / $km^2$). Shrublands had the second highest density (137.8 refugia / $km^2$), followed by low duneveld (102.9 refugia / $km^2$) and high duneveld (58.2 refugia / $km^2$). The pattern was consistent when looking at boltholes and sleeping burrows separately, while more disparate for boltholes. We found that all refugia were distributed disproportionally from habitat availability ($\chi^2$ = 2,303.6; $p < 0.05$). This pattern was consistent for boltholes (2,072.2; $p < 0.05$) and sleeping burrows (297.4; $p < 0.05$). Standardized residuals and selection ratios showed a higher refugia concentration in the riverbed and pans and, to a lesser extent, shrublands. Low duneveld and high duneveld, however, had disproportionally smaller refugia densities compared to available areas (Table S1.8; Figure S1.4).

**Table S1.8.** Distribution of refugia in the study area. SB is sleeping burrows. The area is the cumulative area covered by each habitat within aggregated group-season-year-specific 95% kernel density estimations.

| Type | Metric | Riverbed and pans | High duneveld | Low Duneveld | Low shrublands | $\chi^2$ |
|---|---|---|---|---|---|---|
| | Area (km²) | 17.15 | 22.11 | **52.37** | 10.43 | |
| Refugia | Total (%) | 3746 (31.6%) | 1287 (10.9%) | **5390 (45.4%)** | 1437 (12.1%) | 2303.5 |
| | Density/km² | **218.44** | 58.21 | 102.91 | 137.77 | |
| | Selection ratio | **1.88** | 0.50 | 0.89 | 1.19 | |
| | Std. residual | 43.06 | -28.58 | -12.79 | 6.82 | |
| Boltholes | Total (%) | 2925 (32.8%) | 843 (9.4%) | **4051 (45.4%)** | 1107 (12.4%) | 2072.1 |
| | Density/km² | **170.56** | 38.13 | 77.35 | 106.13 | |
| | Selection ratio | **1.95** | 0.44 | 0.88 | 1.21 | |
| | Std. residual | 40.34 | -28.02 | -11.21 | 6.81 | |
| Burrows | Total (%) | 821 (28%) | 444 (15.1%) | **1339 (45.6%)** | 330 (11.2%) | 297.4 |
| | Density/km² | **47.87** | 20.08 | 25.57 | 31.64 | |
| | Selection ratio | **1.67** | 0.70 | 0.89 | 1.10 | |
| | Std. residual | 16.20 | -8.59 | -6.15 | 1.84 | |

Used habitat (%): Riverbed and pans: 16.8%; High duneveld: 21.7%; Low Duneveld: 51.3%; Low shrublands: 10.2%. Selection ratios > 1 suggest over-representation relative to area, while < 1 suggest under-representation. Std. residuals > 2 suggest over-representation, while < -2 suggest under-representation. $\chi^2$ goodness-of-fit tests were all significant at $p < 0.05$.

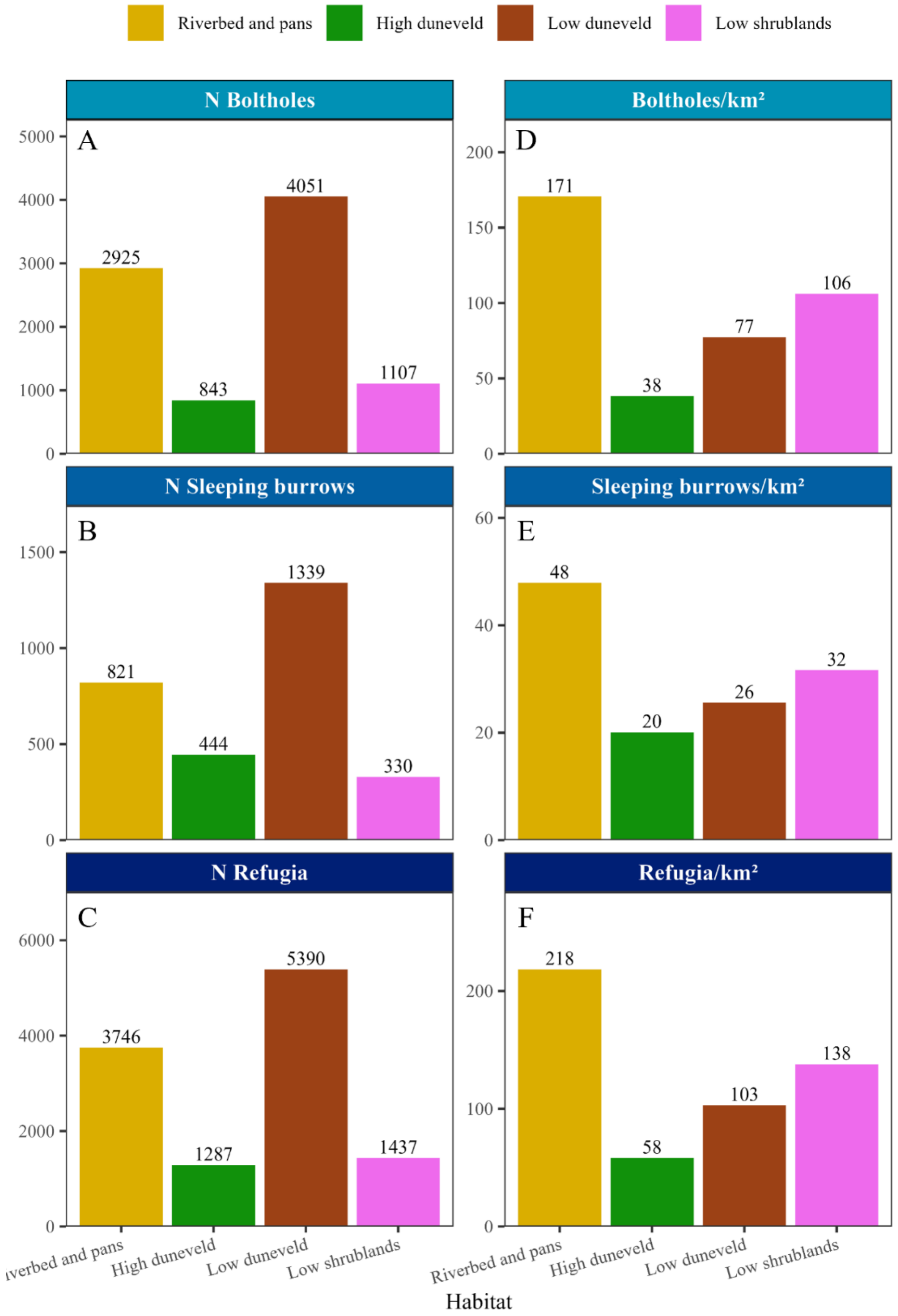

**Figure S1.4.** Visualization of the distribution of refugia across the habitats used by meerkats. (A) Absolute counts of boltholes within each habitat. (B) Absolute counts of sleeping burrows within each habitat. (C) Absolute counts of refugia (i.e. boltholes + sleeping burrows) within each habitat. (D) Bolthole density per habitat. (E) Sleeping burrow density per habitat. (F) Refugia (boltholes and sleeping burrows) density per habitat.

### 6. Alarm-Calling Propensity Analysis

#### *6A - Alarm-Calling Propensity Methods*

We tested how spatial, weather, and social factors influence alarm-calling propensity at each GPS location using generalized linear mixed models (GLMMs) with a binomial structure comparing locations with eagle and vulture alarm calls (1) to all foraging locations without alarm calls in response to any stimulus (0). To account for the breeding season of eagles and vultures affecting abundances and therefore encounter probability, we conducted season-specific analyses (Table S1.9). For the wet season, we compared the 171,807 locations without any alarm to the 13,803 alarm calls. For the dry season, the dataset consisted of 110,266 locations without any alarm and 14,774 alarm calls.

**Table S1.9.** Summary of the raw number of alarm calls in response to the different species. N is the absolute alarm calls count and % is the percentage of all GPS locations recorded per season.

| Threat level | Species | Wet season | | Dry season | | Change |
|---|---|---|---|---|---|---|
| | | N | % | N | % | |
| Non-predator | White-backed vulture | 8,010 | 4.4 | 9,389 | 7.7 | Up |
| | Lappet-faced vulture | 1,539 | 0.8 | 1,955 | 1.6 | Up |
| | Und. vulture | 555 | 0.3 | 549 | 0.4 | Up |
| | Cape vulture | 40 | 0.0 | 67 | 0.1 | Up |
| | White-headed vulture | 12 | 0.0 | 7 | 0.0 | |
| | Palm nut vulture | 0 | 0.0 | 1 | 0.0 | |
| Predator | Martial eagle | 1,965 | 1.1 | 1,737 | 1.4 | Up |
| | **Black-breasted snake eagle** | **1,204** | **0.7** | **525** | **0.4** | **Down** |
| | Tawny eagle | 235 | 0.1 | 215 | 0.2 | Up |
| | Bateleur | 87 | 0.0 | 223 | 0.2 | Up |
| | Und. eagle | 54 | 0.0 | 51 | 0.0 | |
| | Brown snake eagle | 53 | 0.0 | 28 | 0.0 | |
| | Steppe eagle | 22 | 0.0 | 9 | 0.0 | |
| | Wahlberg's eagle | 17 | 0.0 | 15 | 0.0 | |
| | Booted eagle | 10 | 0.0 | 3 | 0.0 | |

We included variables representing contexts anticipated to affect encounter rate, detection capacity, and risk perception as predictors. We included habitat type, NDVI, and Topographic Position Index (TPI) as spatial predictors, visibility conditions, daily maximum temperature, as well as drought conditions as weather predictors, and group size and the presence of pups as social predictors. We further included an interaction between habitat type and NDVI.

We calculated TPI using a 3x3 pixel moving window from a 30-m resolution NASA NASADEM digital elevation model (DEM). Given the topography of the study site, TPI values around zero indicate flat terrain, while positive and negative extremes suggest dune crests and interdunal swales, respectively. Additionally, we extracted daily maximum temperature data from the NOAA Climate Prediction Centre (NOAA 2026a), which covers the full time span of the dataset and is strongly correlated with measurements from the temporally limited on-site weather station (Pearson's r = 0.949; 95% CI: 0.946 - 0.952; Table S1.10; Figure S1.5).

**Table S1.10.** Daily maximum temperature seasonal ranges per year.

| | Wet | | Dry | |
|---|---|---|---|---|
| **Year** | **Low T° (°C)** | **High T° (°C)** | **Low T° (°C)** | **High T° (°C)** |
| 2002 | | | 7.49 | 37.19 |
| 2002-2003 | 15.68 | 41.45 | 8.02 | 33.21 |
| 2003-2004 | 19.58 | 39.73 | 10.00 | 34.32 |
| 2004-2005 | 20.17 | 42.37 | 11.95 | 35.86 |
| 2005-2006 | 18.05 | 41.05 | 6.84 | 36.02 |
| 2006-2007 | 14.08 | 42.40 | 9.65 | 38.08 |
| 2007-2008 | 16.57 | 40.85 | 12.07 | 36.95 |
| 2008-2009 | 23.12 | 41.33 | 13.14 | 36.19 |
| 2009-2010 | 14.98 | 42.32 | 7.25 | 38.11 |
| 2010-2011 | 19.64 | 42.13 | 9.92 | 37.07 |
| 2011-2012 | 18.66 | 42.69 | 5.39 | 37.51 |
| 2012-2013 | 20.68 | 42.60 | 14.03 | 34.73 |
| 2013-2014 | 22.33 | 42.17 | 13.88 | 38.14 |
| 2014-2015 | 20.11 | 42.27 | 12.24 | 39.87 |
| 2015-2016 | 23.49 | 44.08 | 15.49 | 36.36 |
| 2016-2017 | 24.98 | 44.36 | 16.26 | 37.76 |
| 2017-2018 | 16.35 | 41.94 | 13.05 | 35.53 |
| 2018-2019 | 23.72 | 42.95 | 10.79 | 38.55 |
| 2019-2020 | 21.81 | 42.37 | 13.73 | 36.53 |
| 2020-2021 | 23.28 | 42.09 | 12.98 | 37.74 |
| 2021-2022 | 21.30 | 42.94 | 13.94 | 37.47 |
| 2022-2023 | 23.44 | 42.55 | 12.83 | 39.14 |
| 2023-2024 | 27.70 | 44.04 | | |

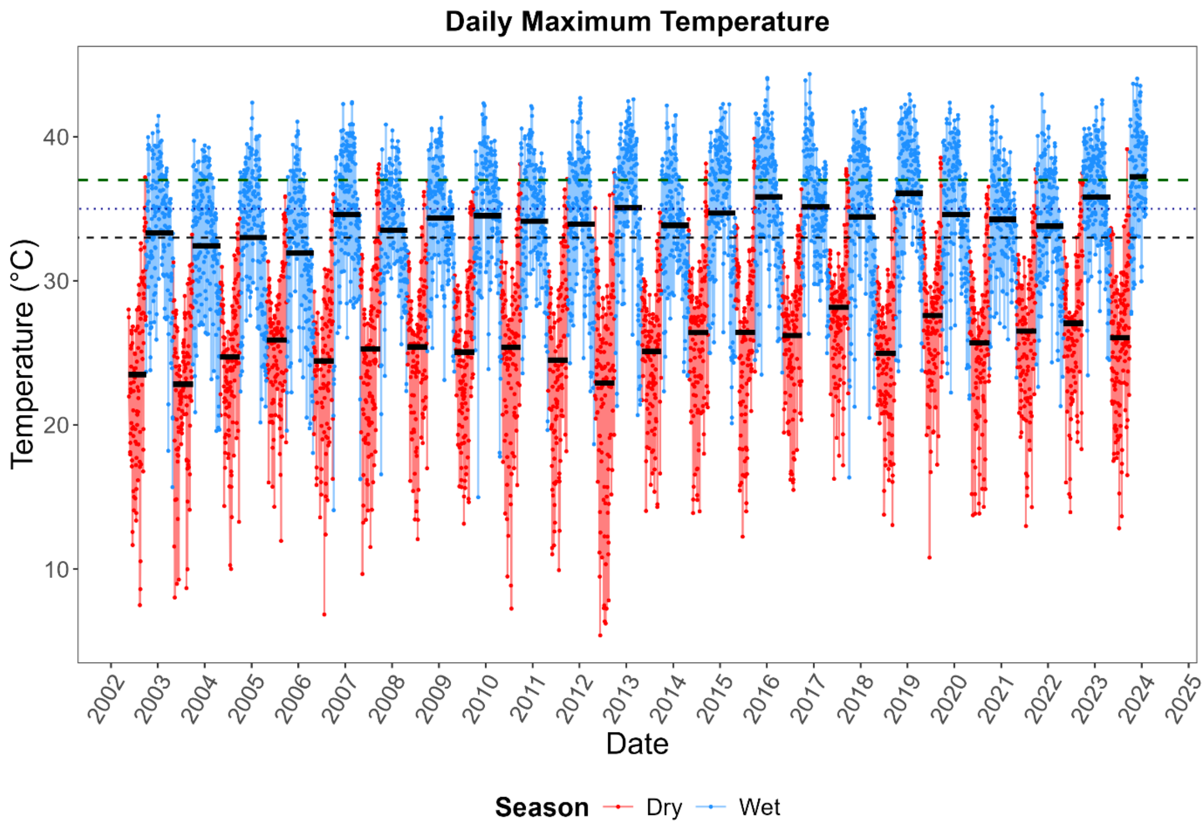

**Figure S1.5.** Daily maximum temperatures over the temporal range of the study. Wet season is blue and dry season is red. The black segments are the seasonal means. The black dashed line is 33°C, the blue dotted line is 35°C, and the green dashed line is 37°C.

Consistent with a previous study at our field site (Groenewoud and Clutton-Brock 2021), we used the *SCI* package version 1.0.3 (Gudmundsson 2025) to calculate the standardised precipitation index (SPI) at 1-, 3-, 6-, and 12-month scales with a gamma distribution based on data from the NOAA Climate Prediction Centre (NOAA 2026b). Doing so, we distinguished periods of severe and extreme droughts (1) from less limiting conditions (0; Figure S1.6).

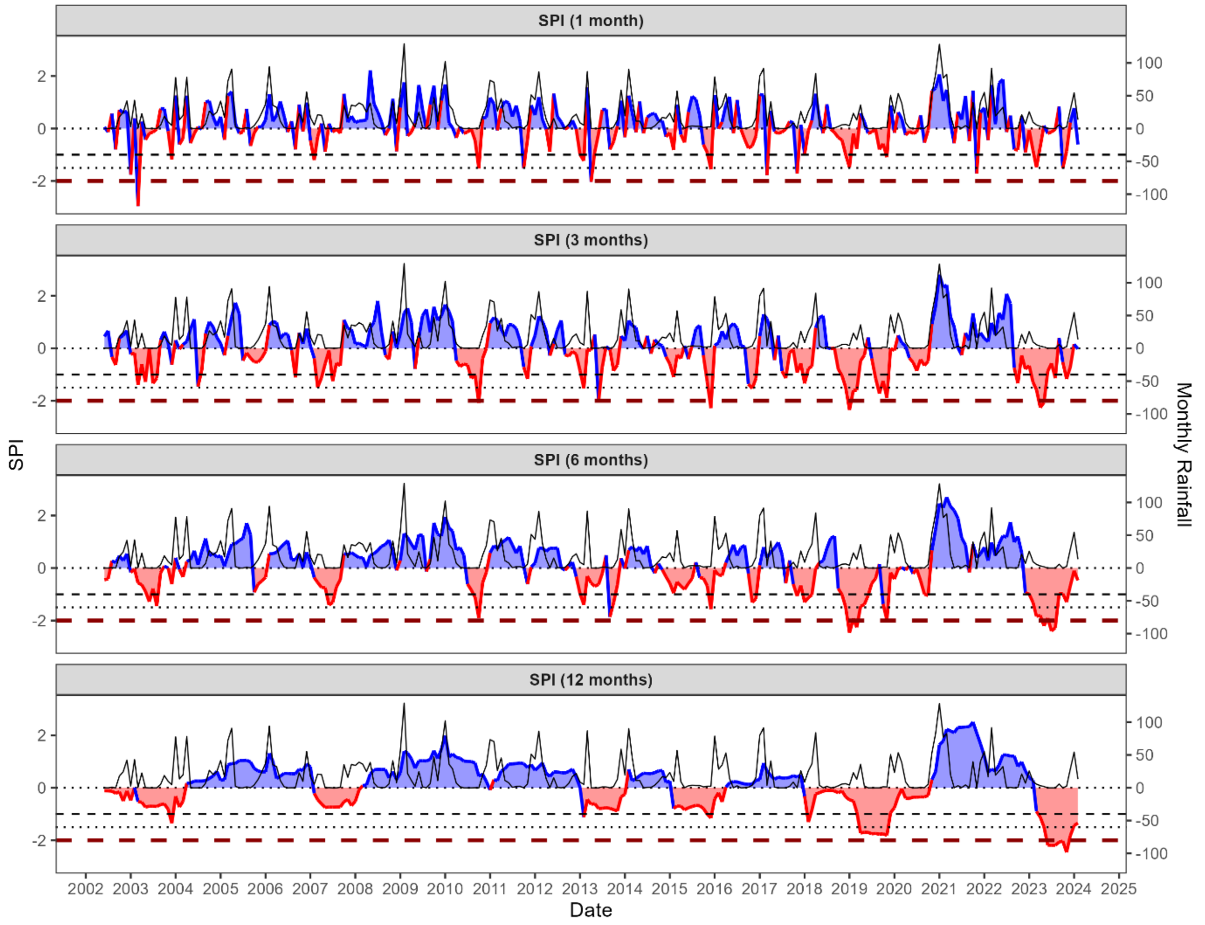

**Figure S1.6.** SPI at different temporal scales. Wet SPI values are shown in blue while dry values are red. The background black line is the monthly rainfall average. The dashed black asymptote is a SPI of -1 delimiting moderate droughts. The dotted black line is a SPI of -1.5 delimiting severe droughts. The red dashed line is a SPI of -2 delimiting extreme droughts.

##### *6B - Preliminary selection steps*

Before proceeding with the alarm-calling propensity analysis, we conducted preliminary selection steps determine the random effects structure and the SPI temporal scale. We fitted a model with the most biologically meaningful structure accounting for the data's temporal structure fitting group nested within follow and year as random intercepts, but it indicated singularity. Therefore, we simplified alternative random effects structures for each SPI temporal scale and compared them by $AIC_c$. This preliminary selection led us to include only group follow as a random intercept for both the wet and dry season analyses of alarm-calling propensity.

**Table S1.11.** Candidate logistic regression models for the meerkat alarm-calling propensity to high-soaring birds between 2002 and 2024 comparing the most complex model structure with drought defined at different temporal scales (1-, 3-, 6-, 12-months) and with different random effect structure. AICc is the corrected Akaike Information Criteria, ΔAICc is the AICc difference with the most supported model, and $w_i$ is the Akaike weight.The most parsimonious models are shown in bold.

| **Dry season** | | | | **Wet season** | | | |
|---|---|---|---|---|---|---|---|
| **Random effects** | **AICc** | **ΔAICc** | **$w_i$** | **Random effects** | **AICc** | **ΔAICc** | **$w_i$** |
| *1-Month* | | | | | | | |
| | | | | **Follow** | **88,464.4** | **0.0** | **0.5** |
| | | | | Group/Follow | 88,466.4 | 2.0 | 0.2 |
| | | | | Group + Follow | 88,466.4 | 2.0 | 0.2 |
| | | | | Year + Group/Follow | 88,468.4 | 4.0 | 0.1 |
| | | | | Year + Group + Follow | 88,468.4 | 4.0 | 0.1 |
| | | | | Year + Group | 95,401.7 | 6,937.3 | 0 |
| | | | | Year | 95,691.0 | 7,226.6 | 0 |
| | | | | Group | 96,003.3 | 7,538.9 | 0 |
| *3-Months* | | | | | | | |
| **Follow** | **83,297.1** | **0** | **0.6** | **Follow** | **88,453.4** | **0.0** | **0.5** |
| Group/Follow | 83,299.1 | 2.0 | 0.2 | Group/Follow | 88,455.4 | 2.0 | 0.2 |
| Group + Follow | 83,299.1 | 2.0 | 0.2 | Group + Follow | 88,455.4 | 2.0 | 0.2 |
| Year + Group/Follow | 83,388.3 | 91.2 | 0 | Year + Group/Follow | 88,457.4 | 4.0 | 0.1 |
| Year + Group + Follow | 83,388.3 | 91.2 | 0 | Year + Group + Follow | 88,457.4 | 4.0 | 0.1 |
| Year + Group | 88,273.9 | 4,976.8 | 0 | Year + Group | 95,351.3 | 6,897.9 | 0 |
| Year | 88,505.1 | 5,208.0 | 0 | Year | 95,640.0 | 7,186.6 | 0 |
| Group | 89,006.5 | 5,709.4 | 0 | Group | 95,969.2 | 7,515.9 | 0 |
| *6-Months* | | | | | | | |
| **Follow** | **83,301.2** | **0** | **0.6** | **Follow** | **88,450.6** | **0.0** | **0.5** |
| Group + Follow | 83,303.2 | 2.0 | 0.2 | Group + Follow | 88,452.6 | 2.0 | 0.2 |
| Group/Follow | 83,303.2 | 2.0 | 0.2 | Group/Follow | 88,452.6 | 2.0 | 0.2 |
| Year + Group + Follow | 83,386.1 | 84.9 | 0 | Year + Group + Follow | 88,454.6 | 4.0 | 0.1 |
| Year + Group/Follow | 83,386.1 | 84.9 | 0 | Year + Group/Follow | 88,454.6 | 4.0 | 0.1 |
| Year + Group | 88,269.2 | 4,968.1 | 0 | Year + Group | 95,348.6 | 6,898.1 | 0 |
| Year | 88,499.9 | 5,198.7 | 0 | Year | 95,635.3 | 7,184.8 | 0 |
| Group | 89,018.5 | 5,717.3 | 0 | Group | 95,972.3 | 7,521.8 | 0 |
| *12-Months* | | | | | | | |
| **Follow** | **83,301.9** | **0** | **0.6** | **Follow** | **88,464.5** | **0.0** | **0.5** |
| Group/Follow | 83,303.9 | 2.0 | 0.2 | Group/Follow | 88,466.5 | 2.0 | 0.2 |
| Group + Follow | 83,303.9 | 2.0 | 0.2 | Group + Follow | 88,466.5 | 2.0 | 0.2 |
| Year + Group + Follow | 83,388.9 | 87.0 | 0 | Year + Group + Follow | 88,468.5 | 4.0 | 0.1 |
| Year + Group/Follow | 83,388.9 | 87.0 | 0 | Year + Group/Follow | 88,468.5 | 4.0 | 0.1 |
| Year + Group | 88,273.1 | 4,971.2 | 0 | Year + Group | 95,406.8 | 6,942.3 | 0 |
| Year | 88,504.8 | 5,203.0 | 0 | Year | 95,694.6 | 7,230.1 | 0 |
| Group | 89,019.0 | 5,718.2 | 0 | Group | 96,009.0 | 7,544.5 | 0 |

We then compared full models with the simplified random effects structure to determine which SPI temporal scale to use in the main alarm-calling propensity analysis. During the dry season, no drought was ever identified using SPI as the 1-month scale. In conclusion, we selected the most parsimonious and used drought at the 3-month scale durng the dry season with follow as random intercept. The pattern was similar for the wet season, but with drought defined at the 6-month scale instead.

**Table S1.12.** Candidate logistic regression models for the meerkat alarm-calling propensity to high-soaring birds between 2002 and 2024 comparing drought defined at different temporal scales (1-, 3-, 6-, 12-months) with group follow as random effect structure. AICc is the corrected Akaike Information Criteria, ΔAICc is the AICc difference with the most supported model, and $w_i$ is the Akaike weight, and SPI is the Standardised Precipitation Index. The most parsimonious models are shown in bold.

| | Dry season | | | Wet season | | |
|---|---|---|---|---|---|---|
| **SPI temporal scale** | **AICc** | **ΔAICc** | **$w_i$** | **AICc** | **ΔAICc** | **$w_i$** |
| **3-Months** | **83,297.12** | **0.00** | **0.82** | 88,453.37 | 2.81 | 0.20 |
| 6-Months | 83,301.17 | 4.05 | 0.11 | **88,450.55** | **0.00** | **0.80** |
| 12-Months | 83,301.85 | 4.73 | 0.08 | 88,464.50 | 13.95 | 0.00 |
| 1-Month | | | | 88,464.41 | 13.85 | 0.00 |

We then proceeded we the alarm-calling propensity analysis and compared 11 candidate models to a null model using $AIC_c$.

##### *6C - Results*

For alarm-calling propensity during the wet and dry seasons, both models with the lowest $AIC_c$ values included TPI, temperature, drought conditions, as well as habitat-specific two-way interactions with NDVI, visibility conditions, group size, and pup presence (Tables S1.13 and S1.14).

Consistently, alarm-calling propensity was lower during the wet rather than dry season (Table S1.15; Figure S1.7). During the wet season, alarm-calling propensity predictions were lower within habitats with a greater refugia density, with $\hat{p}$ = 2.5% (95% CI: 2.3 – 2.8%; Figure S1.7A) for the riverbed and pans and 2.6% (2.2 – 3.1%) for low shrublands compared to 3.2% (2.9 – 3.5%). This baseline habitat structure was similar during the dry season, with high duneveld (9.3%; 7.8 – 11.0%) and low shrublands (7.2%; 6.1 – 8.5%) being marginally higher than low duneveld (8.0%; 7.4 – 8.7%), while alarm-calling propensity was considerably lower within the riverbed and pans (5.4%; 4.9 – 6.0%). During the wet season, alarm-calling propensity marginally increased with topographic exposure from 2.9% (2.5 – 3.4%) to 3.8% (3.0 – 4.8%) over the range of TPI values (Figure S1.7B). This probability followed a similar marginal increase during the dry season, from 7.1% (6.0 – 8.4%) to 9.4% (7.7 – 11.3%). Alarm-calling propensity was more likely with higher vegetation productivity during both seasons (Figure S1.7C-F). From the $2.5^{th}$ to the $97.5^{th}$ season-specific NDVI values, alarm-calling predicted probabilities for the reference low duneveld increased from 2.9% (2.6 – 3.2%) to 4.3% (3.7 – 4.9%) during the wet season and from 7.2% (6.5 – 7.8%) to 9.7% (8.5 – 11.0%) during the dry season (Figure S1.7E). This increase in NDVI only differed from low duneveld within riverbed and pans (Figure S1.7C), with a greater increase during the wet season from 2.1% (1.9 – 2.3%) to 3.9% (3.4 – 4.5%) and a marginally larger increase during the dry season from 4.6% (4.1 – 5.1%) to 7.1% (6.2 – 8.1%).

Temperature had the strongest effects on alarm-calling propensity, which decreased in the wet season from 15.1% (13.0 – 17.4%) at 14.1°C to 1.5% (1.4 – 1.7%) at 44.4°C and from 20.7% (18.5 – 23.1%) at 5.4°C to 4.0% (3.6 – 4.4%) at 39.9°C during the dry season (Figure S1.7G). Alarm-calling propensity decreased slightly under drought conditions from 3.2% (2.9 – 3.5%) to 2.5% (2.1 – 2.9%) during the wet season and from 8.0% (7.4 – 8.7%) to 6.6% (5.5 – 7.8%) during the dry season (Figure S1.7H). We did not detect an effect of poor visibility on alarm-calling propensity in low duneveld during both season (Figure S1.7I). However, poor visibility was associated with weak decreases during the wet season from 2.5% (2.3 – 2.8%) to 1.9% (1.8 – 2.1%) in the riverbed and pans and from 2.6% (2.2 – 3.1%) to 2.0% (1.8 – 2.3%) in the low shrublands. These decreases were larger during the dry season, dropping from 5.4% (4.9 – 6.0%) to 4.4% (4.1 – 4.7%) in the riverbed and pans and from 7.2% (6.1 – 8.5%) to 5.2% (4.6 – 5.7%) in low shrublands.

We found a marginal effect of pup presence on alarm-calling propensity during the wet season in the riverbed and pans, from 2.5% (2.3 – 2.8%) to 2.7% (2.4 –3.0%), but not in other habitats (Figure S1.7J). During the dry season, however, groups with pups had a lower alarm-calling propensity in low duneveld, decreasing from 8.0% (7.4 – 8.7%) to 6.6% (6.0 – 7.4%), which was consistent across habitats. Group size, ranging from 3 to 27 (2.5$^{th}$ to 97.5$^{th}$ percentiles) during the wet season, increased in low duneveld from 2.7% (2.4 – 3.0%) to 4.2% (3.6 – 4.8%; Figure S1.7M), marginally differing with the smaller increases in riverbed and pans from 2.3% (2.0 – 2.5%) to 3.0% (2.6 – 3.4%; Figure S1.7K) and in low shrublands from 2.4% (2.0 – 2.9%) to 3.1% (2.4 – 4.0%; Figure S1.7N). During the dry season, alarm-calling propensity increased in low duneveld from 6.6% (6.0 – 7.3%) for groups of 4 individuals to 11.0% (9.9 – 12.3%; Figure S1.7M) for group of 31 individuals. This increase marginally differed for low shrublands (Figure S1.7N), rising from 6.3% (5.2 – 7.7%) to 9.0% (7.1 – 11.2%).

**Table S1.13.** Candidate logistic regression models for the meerkat alarm-calling propensity to high-soaring birds during the wet season between 2002 and 2024. K is the degrees of freedom, $AIC_c$ is the corrected Akaike Information Criteria, $w_i$ is the Akaike weight, and $\Delta AIC_c$ is the $AIC_c$ difference with the most supported model, and T° is the daily maximum temperature. The most parsimonious model is shown in bold.

| Model | K | AICc | ΔAICc | $w_i$ |
|---|---|---|---|---|
| **Habitat + NDVI + Habitat:NDVI + TPI + Visibility + T° + Drought + Group size + Pups + Habitat:Visibility + Habitat:Group size + Habitat:Pups** | **24** | **88,435.55** | **0** | **0.94** |
| Habitat + NDVI + Habitat:NDVI + TPI + Visibility + T° + Drought + Group size + Pups | 15 | 88,440.97 | 5.42 | 0.06 |
| Habitat + NDVI + Habitat:NDVI + TPI + Visibility + T° + Drought + Habitat:Visibility | 16 | 88,458.84 | 23.28 | 0 |
| Habitat + NDVI + Habitat:NDVI + TPI + Visibility + T° + Drought | 13 | 88,470.02 | 34.46 | 0 |
| Visibility + T° + Drought + Group size + Pups | 7 | 88,747.97 | 312.41 | 0 |
| Visibility + T° + Drought | 5 | 88,776.52 | 340.96 | 0 |
| Habitat + NDVI + Habitat:NDVI + TPI + Group size + Pups | 12 | 88,980.55 | 545 | 0 |
| Habitat + NDVI + Habitat:NDVI + TPI + Group size + Pups + Habitat:Group size + Habitat:Pups | 18 | 88,985.75 | 550.20 | 0 |
| Habitat + NDVI + Habitat:NDVI + TPI | 10 | 89,021.77 | 586.22 | 0 |
| Group size + Pups | 4 | 89,342.95 | 907.40 | 0 |
| Null | 2 | 89,389.20 | 953.65 | 0 |

**Table S1.14.** Candidate logistic regression models for the meerkat alarm-calling propensity to high-soaring birds during the dry season between 2002 and 2024. K is the degrees of freedom, $AIC_c$ is the corrected Akaike Information Criteria, $w_i$ is the Akaike weight, and $\Delta AIC_c$ is the $AIC_c$ difference with the most supported model, and T° is the daily maximum temperature. The most parsimonious model is shown in bold.

| Model | K | AICc | ΔAICc | $w_i$ |
|---|---|---|---|---|
| **Habitat + NDVI + Habitat:NDVI + TPI + Visibility + T° + Drought + Group size + Pups + Habitat:Visibility + Habitat:Group size + Habitat:Pups** | **24** | **83,297.12** | **0** | **0.79** |
| Habitat + NDVI + Habitat:NDVI + TPI + Visibility + T° + Drought + Group size + Pups | 15 | 83,299.75 | 2.63 | 0.21 |
| Habitat + NDVI + Habitat:NDVI + TPI + Visibility + T° + Drought + Habitat:Visibility | 16 | 83,411.97 | 114.85 | 0 |
| Habitat + NDVI + Habitat:NDVI + TPI + Visibility + T° + Drought | 13 | 83,422.21 | 125.09 | 0 |
| Habitat + NDVI + Habitat:NDVI + TPI + Group size + Pups | 12 | 83,666.73 | 369.61 | 0 |
| Habitat + NDVI + Habitat:NDVI + TPI + Group size + Pups + Habitat:Group size + Habitat:Pups | 18 | 83,673.05 | 375.93 | 0 |
| Visibility + T° + Drought + Group size + Pups | 7 | 83,787.48 | 490.36 | 0 |
| Habitat + NDVI + Habitat:NDVI + TPI | 10 | 83,823.03 | 525.91 | 0 |
| Visibility + T° + Drought | 5 | 83,879.47 | 582.35 | 0 |
| Group size + Pups | 4 | 84,137.02 | 839.90 | 0 |
| Null | 2 | 84,258.94 | 961.82 | 0 |

**Table S1.15.** Estimates from the most parsimonious models for meerkat alarm-calling propensity in response to high-soaring birds during the wet and dry seasons. The seasonal models were fitted with 185,610 observations from 28,627 follows during the wet season and 125,040 observations from 20,115 follows during the dry season. Confidence intervals excluding 0 are shown in bold and underlined for the wet and dry season respectively. Abbreviations: SE, standard error; CI, confidence interval; $\sigma^2$, variance of random effects; NDVI, Normalised Difference Vegetation Index; TPI, Topographic Position Index. Reference levels are low duneveld for habitat type, fine visibility, no drought, and no pups.

| | **Wet season** | | | **Dry season** | | |
|---|---|---|---|---|---|---|
| **Term** | **Estimate** | **SE** | **95% CI** | **Estimate** | **SE** | **95% CI** |
| **Fixed effects** | | | | | | |
| Intercept | **-3.407** | **0.051** | **-3.51 – -3.31** | **-2.437** | **0.043** | **-2.52 – -2.35** |
| *Spatial effects* | | | | | | |
| Riverbed and pans | **-0.254** | **0.061** | **-0.37 – -0.13** | **-0.422** | **0.060** | **-0.54 – -0.30** |
| High duneveld | 0.120 | 0.110 | -0.10 – 0.33 | 0.161 | 0.101 | -0.04 – 0.36 |
| Low shrublands | **-0.202** | **0.09** | **-0.38 – -0.03** | -0.120 | 0.094 | -0.30 – 0.06 |
| TPI | 0.017 | 0.010 | 0 – 0.04 | 0.017 | 0.010 | 0 – 0.04 |
| NDVI | **0.110** | **0.021** | **0.07 – 0.15** | **0.088** | **0.023** | **0.04 – 0.13** |
| Riverbed and pans × NDVI | **0.061** | **0.027** | **0.01 – 0.11** | 0.039 | 0.029 | -0.02 – 0.09 |
| High duneveld × NDVI | -0.017 | 0.074 | -0.16 – 0.13 | -0.006 | 0.067 | -0.14 – 0.13 |
| Low shrublands × NDVI | 0.002 | 0.042 | -0.08 – 0.08 | 0.018 | 0.048 | -0.08 – 0.11 |
| *Weather effects* | | | | | | |
| Poor visibility | -0.031 | 0.050 | -0.13 – 0.07 | -0.021 | 0.046 | -0.11 – 0.07 |
| Temperature | **-0.350** | **0.015** | **-0.38 – -0.32** | **-0.287** | **0.015** | **-0.32 – -0.26** |
| Drought | **-0.277** | **0.074** | **-0.42 – -0.13** | **-0.220** | **0.092** | **-0.40 – -0.04** |
| *Social effects* | | | | | | |
| Group size | **0.111** | **0.020** | **0.07 – 0.15** | **0.144** | **0.019** | **0.11 – 0.18** |
| Pups | -0.035 | 0.043 | -0.12 – 0.05 | **-0.207** | **0.050** | **-0.30 – -0.11** |
| *Context-dependence* | | | | | | |
| Riverbed and pans × poor visibility | **-0.250** | **0.066** | **-0.38 – -0.12** | **-0.209** | **0.066** | **-0.34 – -0.08** |
| High duneveld × poor visibility | -0.069 | 0.130 | -0.32 – 0.19 | -0.156 | 0.114 | -0.38 – 0.07 |
| Low shrublands × poor visibility | **-0.247** | **0.099** | **-0.44 – -0.05** | **-0.335** | **0.101** | **-0.53 – -0.14** |
| Riverbed and pans × group size | -0.041 | 0.028 | -0.10 – 0.01 | 0.002 | 0.027 | -0.05 – 0.05 |
| High duneveld × group size | -0.042 | 0.062 | -0.16 – 0.08 | -0.057 | 0.066 | -0.19 – 0.07 |
| Low shrublands × group size | -0.048 | 0.042 | -0.13 – 0.03 | -0.046 | 0.041 | -0.13 – 0.03 |
| Riverbed and pans × pups | 0.112 | 0.059 | 0 – 0.23 | 0.017 | 0.070 | -0.12 – 0.15 |
| High duneveld × pups | 0.048 | 0.123 | -0.19 – 0.29 | -0.128 | 0.130 | -0.38 – 0.13 |
| Low shrublands × pups | -0.024 | 0.089 | -0.20 – 0.15 | 0.076 | 0.103 | -0.13 – 0.28 |
| **Random effects** | **$\sigma^2$** | | | **$\sigma^2$** | | |
| Group follow | 2.7 | | | 1.8 | | |

**Table S1.16.** Odds ratios from the most parsimonious models for alarm-calling propensity to high-soaring birds during the wet and dry seasons. 95% confidence intervals excluding 1 are shown in bold and underlined for the wet and dry season respectively. Abbreviations: OR, odds ratios; CI, confidence interval; $\sigma^2$, variance; NDVI, Normalised Difference Vegetation Index; TPI, Topographic Position Index.

| | **Wet season** | | **Dry season** | |
|---|---|---|---|---|
| **Term** | **OR** | **95% CI** | **OR** | **95% CI** |
| **Intercept** | **0.033** | **0.03 – 0.04** | 0.087 | 0.08 – 0.10 |
| *Spatial effects* | | | | |
| **Riverbed and pans** | **0.776** | **0.69 – 0.87** | 0.656 | 0.58 – 0.74 |
| High duneveld | 1.127 | 0.91 – 1.40 | 1.174 | 0.96 – 1.43 |
| **Low shrublands** | **0.817** | **0.68 – 0.98** | 0.887 | 0.74 – 1.07 |
| TPI | 1.017 | 1.00 – 1.04 | 1.017 | 1.00 – 1.04 |
| **NDVI** | **1.116** | **1.07 – 1.16** | 1.092 | 1.04 – 1.14 |
| **Riverbed and pans × NDVI** | **1.063** | **1.01 – 1.12** | 1.039 | 0.98 – 1.10 |
| High duneveld × NDVI | 0.983 | 0.85 – 1.14 | 0.994 | 0.87 – 1.13 |
| Low shrublands × NDVI | 1.002 | 0.92 – 1.09 | 1.018 | 0.93 – 1.12 |
| *Weather effects* | | | | |
| Poor visibility | 0.969 | 0.88 – 1.07 | 0.979 | 0.90 – 1.07 |
| **Temperature** | **0.704** | **0.68 – 0.73** | 0.751 | 0.73 – 0.77 |
| **Drought** | **0.758** | **0.66 – 0.88** | 0.802 | 0.67 – 0.96 |
| *Social effects* | | | | |
| **Group size** | **1.117** | **1.07 – 1.16** | 1.155 | 1.11 – 1.20 |
| Pups | 0.965 | 0.89 – 1.05 | 0.813 | 0.74 – 0.90 |
| *Context-dependence* | | | | |
| **Riverbed and pans × poor visibility** | **0.779** | **0.68 – 0.89** | 0.811 | 0.71 – 0.92 |
| High duneveld × poor visibility | 0.933 | 0.72 – 1.20 | 0.856 | 0.68 – 1.07 |
| **Low shrublands × poor visibility** | **0.781** | **0.64 – 0.95** | 0.715 | 0.59 – 0.87 |
| Riverbed and pans × group size | 0.96 | 0.91 – 1.01 | 1.002 | 0.95 – 1.06 |
| High duneveld × group size | 0.959 | 0.85 – 1.08 | 0.945 | 0.83 – 1.08 |
| Low shrublands × group size | 0.954 | 0.88 – 1.04 | 0.955 | 0.88 – 1.04 |
| Riverbed and pans × pups | 1.119 | 1.00 – 1.26 | 1.017 | 0.89 – 1.17 |
| High duneveld × pups | 1.049 | 0.82 – 1.33 | 0.880 | 0.68 – 1.14 |
| Low shrublands × pups | 0.976 | 0.82 – 1.16 | 1.079 | 0.88 – 1.32 |

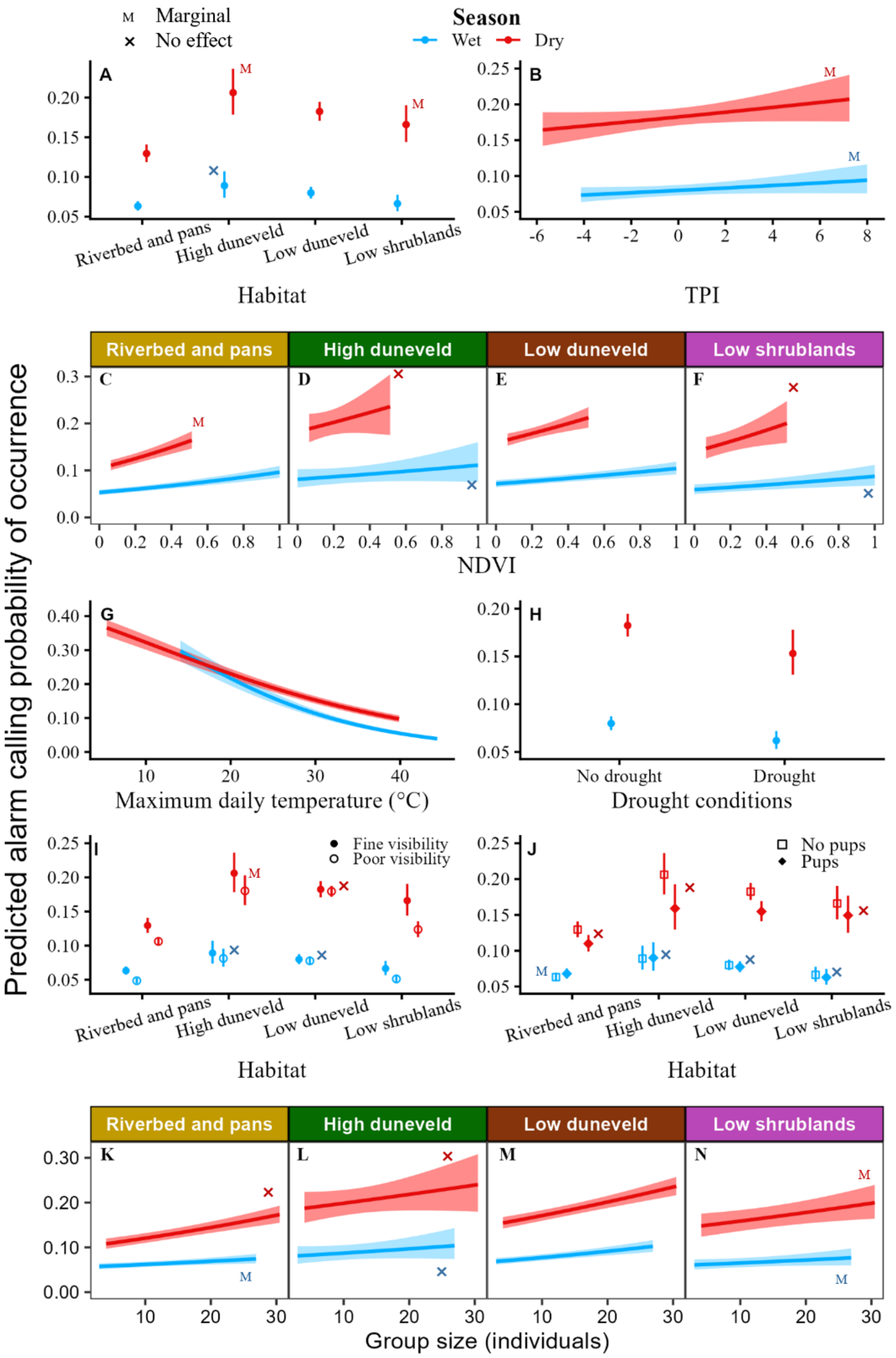

**Figure S1.7.** Predicted values for all the effects of the final model of the objective on alarm-calling propensity for the wet (blue; marginal $\delta R^2$ : 0.04; conditional $\delta R^2$ : 0.47) and dry (red; marginal $\delta R^2$ : 0.04; conditional $\delta R^2$ : 0.38) seasons for which the 95% confidence intervals did not include 0. Error bars and ribbons are the 95% confidence intervals. To observe the effect better, y axes were scaled accordingly. (A) Baseline habitat structure. (B) Topographic Position Index. (C, D, E, F) habitat-specific Normalised Difference Vegetation Index (NDVI) effects predicted over a more realistic NDVI range excluding 5% of extreme values, which was scaled to range between 0 and 1. (G) Maximum daily temperature. (H) Drought conditions. (I) Habitat-specific visibility effects. (J) Habitat-specific effects of pups. (K, L, M, N) Habitat-specific group size effects predicted over a more realistic group size range excluding 5% of extreme values.

##### *6D - Alarm-Calling Propensity Interpretation*

In our study site high soaring species (eagles and vultures) breed during the dry season, increasing their abundance and activity (Maclean 1969; Ferguson-Lees 2005), a pattern previously observed in vultures at our field site (Thorley and Clutton-Brock 2017). This likely explains the consistently higher alarm-calling propensity during the dry season. All other spatial and weather predictors accounting for detection capacity on alarm-calling propensity aligned with our predictions, particularly the large temperature effects. Endotherms avoid thermoregulatory costs by adjusting activity patterns under high temperatures (Du Plessis et al. 2012; Pattinson et al. 2020; Schoenjahn et al. 2022), including reducing hunting effort in raptors (Austin and Miller 1982). Meerkats similarly increase time under shade as a heat-avoidance tactic (Habicher 2009), likely lowering predator detection capacity. This pattern is consistent with temperature-driven decreases in vigilance and activity across taxa (Du Plessis et al. 2012; Rabaiotti and Woodroffe 2019). We further found no effect of poor visibility on threat discrimination. This suggests that the alarm-calling propensity decrease in shrublands with poor visibility conditions may reflect a lesser detection capacity due to amplified visual obstruction in thicker vegetation (Camp et al. 2012) rather than uncertainty (Hayes and Huntly 2005; Beauchamp 2010). As expected, topographic exposure increased alarm-calling propensity, although weakly, presumably due to better sightlines (Embar et al. 2011). Together, these results support that alarm-calling propensity is a compound process including at least encounter probability and meerkat detection capacity. However, no mechanistic interpretation can be inferred by simply looking at alarm-calling propensity results concerning visibility, group size, and pup presence whose effects may be due to either detection capacity or risk perception, or more likely, a combination of both.

The effects of habitat and vegetation productivity suggest accessibility to refugia may influence meerkat proactive responses to predation risk. We showed that boltholes and sleeping burrows are not distributed evenly across habitat-types, but rather more densely located in the riverbed and pans and shrublands, whereas they are sparse in low duneveld and scarce in high duneveld (see Refugia Analysis). Our alarm-calling propensity results for closely mirrors this pattern, particularly for open pans. Thorley et al. (2026) found that meerkats' sleeping burrow selection also follows this distribution and suggested that substrate properties allowed for more solid structures, thereby facilitating digging. While our results could suggest a lower perceived risk with refugia density, we cannot make such conclusion as alarm-calling propensity represents a compound process. Our findings align with Belding's ground squirrels (*Spermophilus beldingi*), which are more likely to emit alarm calls when they are in a vulnerable location, such as being far from a burrow (Sherman 1985). Likewise, Eastern gray squirrels (*Sciurus carolinensis*) exhibit a lower flight threshold to predators with increasing distance to the nearest refuge tree (Dill and Houtman 1989).

The observed weak increases in alarm-calling propensity with NDVI across habitats and seasons suggest meerkats do not benefit from vegetation for concealment, contrary to our expectations. Instead, thicker vegetation may impair meerkat visibility, thereby detection capacity, resulting in a higher uncertainty (Lagory 1986; van der Merwe and Brown 2008; Camp et al. 2012). Brown bears (*Ursus arctos*) illustrate a similar concealment/detection tradeoff as they avoid habitats constraining movement and rather select more for open landscapes that favour mate and competitor detection (Zarzo-Arias et al. 2025). Although our fenced site excludes large terrestrial predators, this sensitivity to obstructed visibility likely evolved under natural conditions where meerkats coexisted with lions (*Panthera leo*), leopards, and cheetahs (*Acinonyx jubatus*), as in the nearby Kalahari Gemsbok National Park (Clutton-Brock et al. 1999).

Our results suggest meerkat social organisation, a component of social systems (Kappeler 2019; Hayes and Schradin 2026), influences detection capacity and risk perception across the landscape. As cooperative breeders, meerkats depend on several social activities related to key vital rates (Ozgul et al. 2014; Clutton-Brock and Manser 2016; Maag et al. 2022; Thorley et al. 2025).Therefore, the observed higher alarm-calling propensity in larger groups likely reflects a greater detection capacity, supporting the many-eyes-hypothesis (Lima 1995; Rieucau and Martin 2008; Beauchamp et al. 2012).

### 7. Additional results

#### *7A - AICc Tables*

Here are the AICc tables for each of the two alarm-calling components.

**Table S1.17.** Candidate logistic regression models for threat discrimination analysis during the wet season between 2002 and 2024. K is the degrees of freedom, $AIC_c$ is the corrected Akaike Information Criteria, $w_i$ is the Akaike weight, and $\Delta AIC_c$ is the $AIC_c$ difference with the most supported model. The most parsimonious model is shown in bold.

| Model | K | $AIC_c$ | $\Delta AIC_c$ | $w_i$ |
|---|---|---|---|---|
| Habitat + NDVI + Habitat:NDVI + Visibility + Group size + Pups | **13** | **15,651.83** | **0** | **0.46** |
| **Habitat + NDVI + Habitat:NDVI + Group size + Pups** | 12 | 15,653.34 | 1.51 | 0.22 |
| Habitat + NDVI + Habitat:NDVI + Visibility | 11 | 15,653.99 | 2.16 | 0.16 |
| Habitat + NDVI + Habitat:NDVI | 10 | 15,655.52 | 3.70 | 0.07 |
| Habitat + NDVI + Habitat:NDVI + Group size + Pups + Habitat:Pups | 15 | 15,655.62 | 3.79 | 0.07 |
| Habitat + NDVI + Habitat:NDVI + Visibility + Habitat:Visibility | 14 | 15,658.79 | 6.96 | 0.01 |
| Habitat + NDVI + Habitat:NDVI + Visibility + Group size + Pups + Habitat:Visibility + Habitat:Pups | 19 | 15,659.07 | 7.24 | 0.01 |
| Visibility + Group size + Pups | 6 | 15,675.64 | 23.81 | 0 |
| Group size + Pups | 5 | 15,677.23 | 25.40 | 0 |
| Visibility | 4 | 15,677.68 | 25.85 | 0 |
| Null | 3 | 15,679.32 | 27.49 | 0 |

**Table S1.18.** Candidate logistic regression models for threat threat discrimination analysis during the dry season between 2002 and 2024. K is the degrees of freedom, $AIC_c$ is the corrected Akaike Information Criteria, $w_i$ is the Akaike weight, and $\Delta AIC_c$ is the $AIC_c$ difference with the most supported model. The most parsimonious model is shown in bold.

| **Model** | **K** | **AICc** | **ΔAICc** | **$w_i$** |
|---|---|---|---|---|
| **Habitat + NDVI + Habitat:NDVI + Group size + Pups** | **12** | **14,065.33** | **0** | **0.59** |
| Habitat + NDVI + Habitat:NDVI + Visibility + Group size + Pups | 13 | 14,067.21 | 1.88 | 0.23 |
| Habitat + NDVI + Habitat:NDVI + Group size + Pups + Habitat:Pups | 15 | 14,067.78 | 2.44 | 0.17 |
| Habitat + NDVI + Habitat:NDVI + Visibility + Group size + Pups + Habitat:Visibility + Habitat:Pups | 19 | 14,074.82 | 9.49 | 0.01 |
| Habitat + NDVI + Habitat:NDVI | 10 | 14,083.79 | 18.46 | 0 |
| Habitat + NDVI + Habitat:NDVI + Visibility | 11 | 14,085.74 | 20.40 | 0 |
| Group size + Pups | 5 | 14,086.59 | 21.26 | 0 |
| Visibility + Group size + Pups | 6 | 14,088.33 | 23 | 0 |
| Habitat + NDVI + Habitat:NDVI + Visibility + Habitat:Visibility | 14 | 14,091.32 | 25.99 | 0 |
| Null | 3 | 14,102.79 | 37.45 | 0 |
| Visibility | 4 | 14,104.67 | 39.34 | 0 |

**Table S1.19.** Candidate logistic regression models for the meerkat response urgency to high-soaring non-predatory birds' alarm calls between 2002 and 2024. K is the degrees of freedom, $AIC_c$ is the corrected Akaike Information Criteria, $w_i$ is the Akaike weight, and $\Delta AIC_c$ is the $AIC_c$ difference with the most supported model. The most parsimonious model is shown in bold.

| **Model** | **K** | **$AIC_c$** | **$\Delta AIC_c$** | **$w_i$** |
|---|---|---|---|---|
| **Habitat + Group size + Pups + Habitat:Pups** | **11** | **19,829.23** | **0** | **0.89** |
| Habitat + Acoustics + Group size + Pups + Habitat:Acoustics + Habitat:Pups | 15 | 19,834.91 | 5.68 | 0.05 |
| Habitat + Group size + Pups | 8 | 19,836.56 | 7.33 | 0.02 |
| Habitat + Acoustics + Group size + Pups | 9 | 19,837.75 | 8.52 | 0.01 |
| Group size + Pups | 5 | 19,837.84 | 8.61 | 0.01 |
| Acoustics + Group size + Pups | 6 | 19,839.04 | 9.81 | 0.01 |
| Habitat | 6 | 19,895.87 | 66.64 | 0 |
| Habitat + Acoustics | 7 | 19,896.76 | 67.53 | 0 |
| Null | 3 | 19,898.50 | 69.27 | 0 |
| Acoustics | 4 | 19,899.40 | 70.17 | 0 |
| Habitat + Acoustics + Habitat:Acoustics | 10 | 19,900.49 | 71.26 | 0 |

**Table S1.20.** Candidate logistic regression models for the meerkat response intensity to high-soaring non-predatory birds' alarm calls between 2002 and 2024. K is the degrees of freedom, $AIC_c$ is the corrected Akaike Information Criteria, $w_i$ is the Akaike weight, and $\Delta AIC_c$ is the $AIC_c$ difference with the most supported model. The most parsimonious model is shown in bold.

| **Model** | **K** | **$AIC_c$** | **$\Delta AIC_c$** | **$w_i$** |
|---|---|---|---|---|
| Habitat + Acoustics + Group size + Pups | 9 | 22,003.40 | 0 | 0.51 |
| **Habitat + Group size + Pups** | **8** | **22,004.65** | **1.25** | **0.28** |
| Habitat + Group size + Pups + Habitat:Pups | 11 | 22,005.46 | 2.06 | 0.18 |
| Habitat + Acoustics + Group size + Pups + Habitat:Acoustics + Habitat:Pups | 15 | 22,009.28 | 5.88 | 0.03 |
| Habitat + Acoustics | 7 | 22,029.64 | 26.24 | 0 |
| Habitat | 6 | 22,030.84 | 27.44 | 0 |
| Habitat + Acoustics + Habitat:Acoustics | 10 | 22,034.76 | 31.36 | 0 |
| Acoustics + Group size + Pups | 6 | 22,041.66 | 38.26 | 0 |
| Group size + Pups | 5 | 22,042.89 | 39.50 | 0 |
| Acoustics | 4 | 22,069.18 | 65.78 | 0 |
| Null | 3 | 22,070.36 | 66.96 | 0 |

#### *7B - Odds Ratios Tables*

**Table S1.21.** Odds ratios from the most parsimonious model for threat discrimination during both seasons. The wet and dry season models were fitted with 13,803 and 14,774 observations, both from 54 groups over 23 years. Confidence intervals excluding 1 are shown in bold for the wet season and underlined for the dry season. Abbreviations: SE, standard error; CI, confidence interval; NDVI, Normalised Difference Vegetation Index. Reference levels are low duneveld for habitat type, and no pups.

| Term | Wet season | | Dry season | |
|---|---|---|---|---|
| | OR | 95% CI | OR | 95% CI |
| **Intercept** | **0.317** | **0.27 – 0.38** | 0.206 | 0.17 – 0.25 |
| *Spatial effects* | | | | |
| **Riverbed and pans** | **1.165** | **1.06 – 1.28** | 1.272 | 1.14 – 1.42 |
| **High duneveld** | **0.714** | **0.57 – 0.9** | 0.961 | 0.77 – 1.2 |
| **Low shrublands** | **1.248** | **1.08 – 1.44** | 1.118 | 0.95 – 1.32 |
| NDVI | 0.936 | 0.87 – 1 | 0.926 | 0.86 – 1 |
| Riverbed and pans × NDVI | 1.053 | 0.97 – 1.15 | 0.97 | 0.88 – 1.07 |
| **High duneveld × NDVI** | **0.746** | **0.56 – 0.99** | 1.228 | 0.99 – 1.52 |
| **Low shrublands × NDVI** | **1.252** | **1.08 – 1.45** | 1.034 | 0.87 – 1.23 |
| *Social effects* | | | | |
| **Group size** | **0.945** | **0.9 – 0.99** | 1.094 | 1.04 – 1.15 |
| Pups | 1.047 | 0.97 – 1.14 | 1.232 | 1.1 – 1.38 |

**Table S1.22.** Odds ratios from the most parsimonious model for the probability of alarm calls eliciting strong reactions. The model was fitted with 21,598 observations from 55 groups over 23 years. Estimate's 95% confidence intervals excluding 0 are shown in bold. Abbreviations: OR, odds ratios; CI, confidence interval; $\sigma^2$, variance; NDVI, Normalised Difference Vegetation Index; TPI, Topographic Position Index. Reference levels are low duneveld for habitat type, and no pups.

| Term | OR | 95% CI | p-value |
|---|---|---|---|
| **Intercept** | **4.331** | **2.87 – 6.53** | **<0.001** |
| *Spatial effects* | | | |
| Riverbed and pans | 1.047 | 0.95 – 1.16 | 0.369 |
| High duneveld | 1.071 | 0.88 – 1.31 | 0.498 |
| Low shrublands | 0.993 | 0.85 – 1.16 | 0.923 |
| *Social effects* | | | |
| **Group size** | **1.105** | **1.06 – 1.15** | **<0.001** |
| **Pups** | **1.184** | **1.06 – 1.32** | **0.002** |
| *Context-dependence* | | | |
| **Riverbed and pans × pups** | **1.28** | **1.08 – 1.52** | **0.005** |
| High duneveld × pups | 0.847 | 0.6 – 1.2 | 0.349 |
| **Low shrublands × pups** | **1.414** | **1.05 – 1.9** | **0.022** |

**Table S1.23.** Odds ratios from the most parsimonious model for the probability of alarm calls eliciting intense reactions. The model was fitted with 17,063 observations from 55 groups over 23 years. Confidence intervals excluding 1 are shown in bold. Abbreviations: OR, odds ratio; CI, confidence interval. Reference levels are low duneveld for habitat type, and no pups.

| Term | OR | 95% CI |
|---|---|---|
| **Intercept** | **1.626** | **1.4 – 1.89** |
| *Spatial effects* | | |
| **Riverbed and pans** | **1.285** | **1.19 – 1.39** |
| **High duneveld** | **1.273** | **1.1 – 1.48** |
| Low shrublands | 1.086 | 0.96 – 1.22 |
| *Social effects* | | |
| **Group size** | **1.117** | **1.07 – 1.16** |
| Pups | 0.976 | 0.91 – 1.05 |

##### *7C - Alternative models*

As mentioned in the main text, two candidate models could not be differentiated with their $AIC_c$ values for both the threat assessment analyses and the response intensity analysis. We therefore selected the most parsimonious models but we report here the results from the alternative models with estimates tables, as well as visualizing the effects (threat assessment: Table S1.24 and Figure S1.8; Response intensity: Table S1.25 and Figure S1.9). The threat discrimination alternative model included habitat in interaction with NDVI, visibility conditions, group size and pup presence as predictors. The response intensity alternative model included habitat, acoustic conditions, group size and pup presence as predictors.

**Table S1.24.** Estimates from the alternative models for threat discrimination during both seasons. The wet and dry season models were fitted with 13,803 and 14,774 observations respectively, both from 54 meerkat groups over 23 years. Confidence intervals excluding 0 are shown in bold for the wet season and underlined for the dry season. Abbreviations: SE, standard error; CI, confidence interval; σ², variance of random effects; NDVI, Normalised Difference Vegetation Index. Reference levels are low duneveld for habitat type, fine visibility, and no pups.

| | Wet season | | | Dry season | | |
|---|---|---|---|---|---|---|
| **Term** | **Estimate** | **SE** | **95% CI** | **Estimate** | **SE** | **95% CI** |
| **Fixed effects** | | | | | | |
| **Intercept** | **-1.04** | **0.108** | **-1.25 – -0.83** | -1.553 | 0.124 | -1.8 – -1.31 |
| *Spatial effects* | | | | | | |
| **Riverbed and pans** | **0.154** | **0.049** | **0.06 – 0.25** | 0.241 | 0.055 | 0.13 – 0.35 |
| **High duneveld** | **-0.337** | **0.116** | **-0.56 – -0.11** | -0.04 | 0.114 | -0.26 – 0.18 |
| **Low shrublands** | **0.219** | **0.073** | **0.08 – 0.36** | 0.112 | 0.085 | -0.06 – 0.28 |
| NDVI | -0.066 | 0.036 | -0.14 – 0 | -0.076 | 0.039 | -0.15 – 0 |
| Riverbed and pans × NDVI | 0.052 | 0.044 | -0.03 – 0.14 | -0.03 | 0.049 | -0.13 – 0.07 |
| **High duneveld × NDVI** | **-0.298** | **0.144** | **-0.58 – -0.02** | 0.206 | 0.108 | -0.01 – 0.42 |
| **Low shrublands × NDVI** | **0.223** | **0.074** | **0.08 – 0.37** | 0.035 | 0.087 | -0.14 – 0.21 |
| *Weather effects* | | | | | | |
| Poor visibility | -0.157 | 0.085 | -0.32 – 0.01 | -0.038 | 0.109 | -0.25 – 0.18 |
| *Social effects* | | | | | | |
| **Group size** | **-0.056** | **0.026** | **-0.11 – -0.01** | 0.09 | 0.027 | 0.04 – 0.14 |
| Pups | 0.047 | 0.042 | -0.03 – 0.13 | 0.21 | 0.057 | 0.1 – 0.32 |
| **Random effects** | **σ²** | | | **σ²** | | |
| Year | 0.126 | | | 0.129 | | |
| Group identity | 0.047 | | | 0.079 | | |

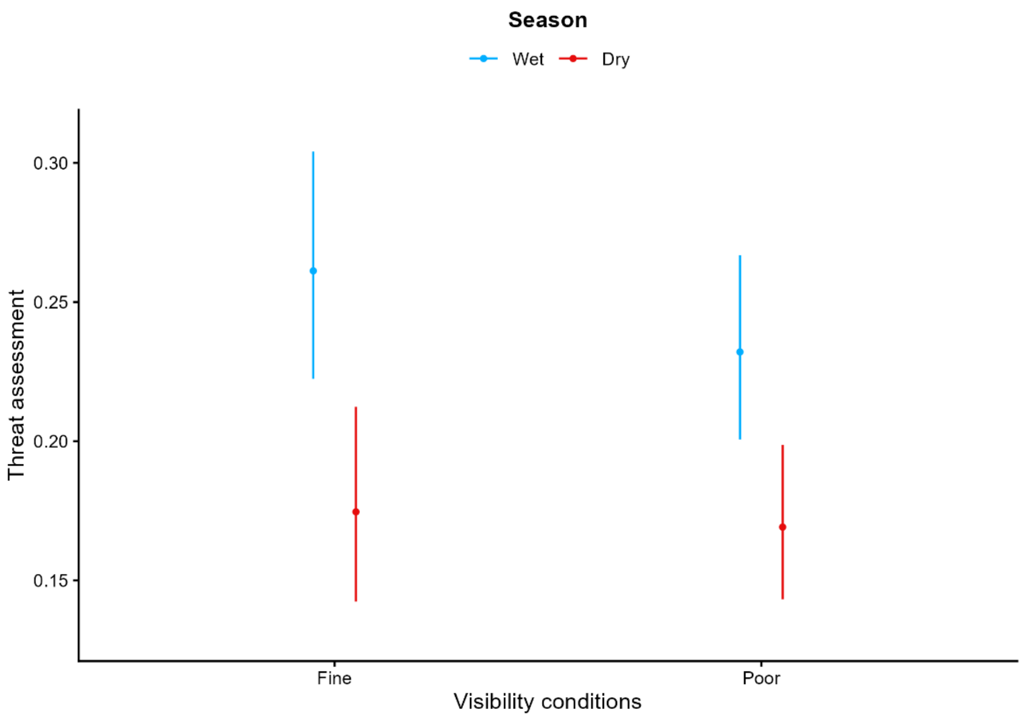

**Figure S1.8.** Predicted values for the visibility effect of the alternative model of the objective on threat discrimination for the wet (blue) and dry (red) seasons. Error bars are the 95% confidence intervals. The visibility effect in the wet season model is marginal, but we did not detect an effect in the dry season model. To observe the effect better, y axes were scaled accordingly.

**Table S1.25.** Estimates from the alternative model for the probability of alarm calls eliciting intense reactions (fleeing vs bipedal vigilance). Confidence intervals excluding 0 are shown in bold. Abbreviations: SE, standard error; CI, confidence interval. Reference levels are low duneveld for habitat type, fine acoustics, and no pups.

| Term | Estimate | SE | 95% CI |
|---|---|---|---|
| **Fixed effects** | | | |
| **Intercept** | **0.47** | **0.078** | **0.32 – 0.62** |
| *Spatial effects* | | | |
| **Riverbed and pans** | **0.251** | **0.041** | **0.17 – 0.33** |
| **High duneveld** | **0.236** | **0.076** | **0.09 – 0.39** |
| Low shrublands | 0.08 | 0.061 | -0.04 – 0.2 |
| *Weather effects* | | | |
| Poor acoustics | 0.077 | 0.043 | -0.01 – 0.16 |
| *Social effects* | | | |
| **Group size** | **0.11** | **0.021** | **0.07 – 0.15** |
| Pups | -0.025 | 0.035 | -0.09 – 0.04 |
| **Random effects** | **$\sigma^2$** | | |
| Year | 0.107 | | |
| Group identity | 0.024 | | |

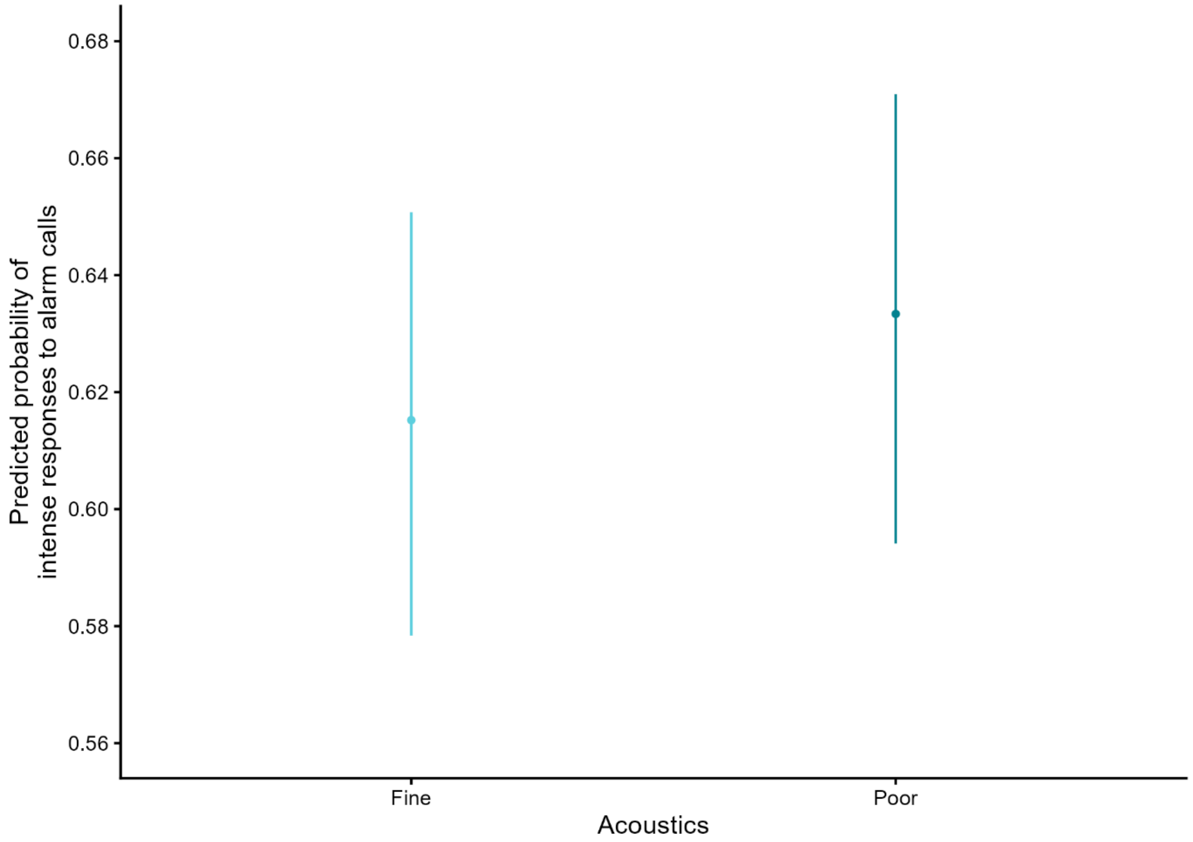

**Figure S1.9.** Predicted values for the marginal effect of acoustic conditions from the alternative model for response escalation with reaction intensity. Error bars are the 95% confidence interval. To observe the effect better, y axes were scaled accordingly.

#### 8. Sensitivity Analyses

Although we restricted our threat discrimination analysis to alarms emitted at eagles and vulture, as separating the wet and dry seasons, there may be season-specific variation in abundances. Therefore, we verified whether adding 'month', and 'fortnight' to the threat discrimination final models for both the wet and dry seasons would generate similar results or large differences. Additionally, as most tall trees are located in the riverbed and pans, we ran the final threat discrimination excluding this habitat.

In conclusion, threat discrimination was different in February, March, April, and October from the January reference. In contrast, there was no evidence of a monthly effect during the dry season. During the wet season, other coefficient remained similar as our main results, but the NDVI effect in low duneveld weakened, and group size weakened by almost half with the 95% confidence interval now overlapping 0 (Table S1.26; Figure S1.10). Thus, we could not consider the wet season group size effect as robust. The 'fortnight' sensitivity analysis indicated weak statistical differences over time, but it is consistent with the monthly pattern (Table S1.27; Figure S1.11). When excluding the riverbed and pans from the dataset, the baseline habitat structure did not change (Table S1.28; Figure S1.12). The NDVI effect slightly weakened, the wet season group size effect changed sign with the 95% confidence interval overlapping '0', and the dry season group size effect weakened, while the pup effects held.

**Table S1.26.** Estimates from threat discrimination final seasonal models including "**Month**" as predictor. Confidence intervals excluding 0 are shown in bold. Abbreviations: SE, standard error; CI, confidence interval; $\sigma^2$, variance of random effects; NDVI, Normalised Difference Vegetation Index. Reference levels are low duneveld for habitat type, fine visibility, and no pups.

| | **Wet season** | | | **Dry season** | | |
|---|---|---|---|---|---|---|
| **Term** | **Estimate** | **SE** | **95% CI** | **Estimate** | **SE** | **95% CI** |
| **Fixed effects** | | | | | | |
| **Intercept** | **-0.974** | **0.1** | **-1.17 – -0.78** | **-1.658** | **0.114** | **-1.88 – -1.44** |
| *Temporal effects* | | | | | | |
| **February** / June | **-0.208** | **0.08** | **-0.37 – -0.05** | 0.074 | 0.074 | -0.07 – 0.22 |
| **March** / July | **-0.242** | **0.076** | **-0.39 – -0.09** | 0.048 | 0.079 | -0.11 – 0.2 |
| **April** / August | **-0.466** | **0.076** | **-0.62 – -0.32** | 0.127 | 0.085 | -0.04 – 0.29 |
| **October** / September | **-0.314** | **0.079** | **-0.47 – -0.16** | 0.117 | 0.091 | -0.06 – 0.3 |
| November | 0.001 | 0.077 | -0.15 – 0.15 | | | |
| December | 0.05 | 0.08 | -0.11 – 0.21 | | | |
| *Spatial effects* | | | | | | |
| **Riverbed and pans** | **0.161** | **0.049** | **0.06 – 0.26** | **0.243** | **0.055** | **0.13 – 0.35** |
| **High duneveld** | **-0.326** | **0.116** | **-0.55 – -0.1** | -0.035 | 0.114 | -0.26 – 0.19 |
| **Low shrublands** | **0.23** | **0.073** | **0.09 – 0.37** | 0.121 | 0.086 | -0.05 – 0.29 |
| NDVI | -0.024 | 0.04 | -0.1 – 0.05 | -0.044 | 0.045 | -0.13 – 0.04 |
| Riverbed and pans × NDVI | 0.05 | 0.043 | -0.03 – 0.14 | -0.04 | 0.05 | -0.14 – 0.06 |
| **High duneveld × NDVI** | **-0.292** | **0.143** | **-0.57 – -0.01** | 0.204 | 0.108 | -0.01 – 0.42 |
| **Low shrublands × NDVI** | **0.228** | **0.074** | **0.08 – 0.37** | 0.032 | 0.087 | -0.14 – 0.2 |
| *Social effects* | | | | | | |
| **Group size** | -0.033 | 0.026 | -0.08 – 0.02 | **0.09** | **0.028** | **0.04 – 0.14** |
| **Pups** | 0.057 | 0.042 | -0.02 – 0.14 | **0.221** | **0.058** | **0.11 – 0.33** |
| **Random effects** | **$\sigma^2$** | | | **$\sigma^2$** | | |
| Year | 0.105 | | | 0.128 | | |
| Group identity | 0.046 | | | 0.082 | | |

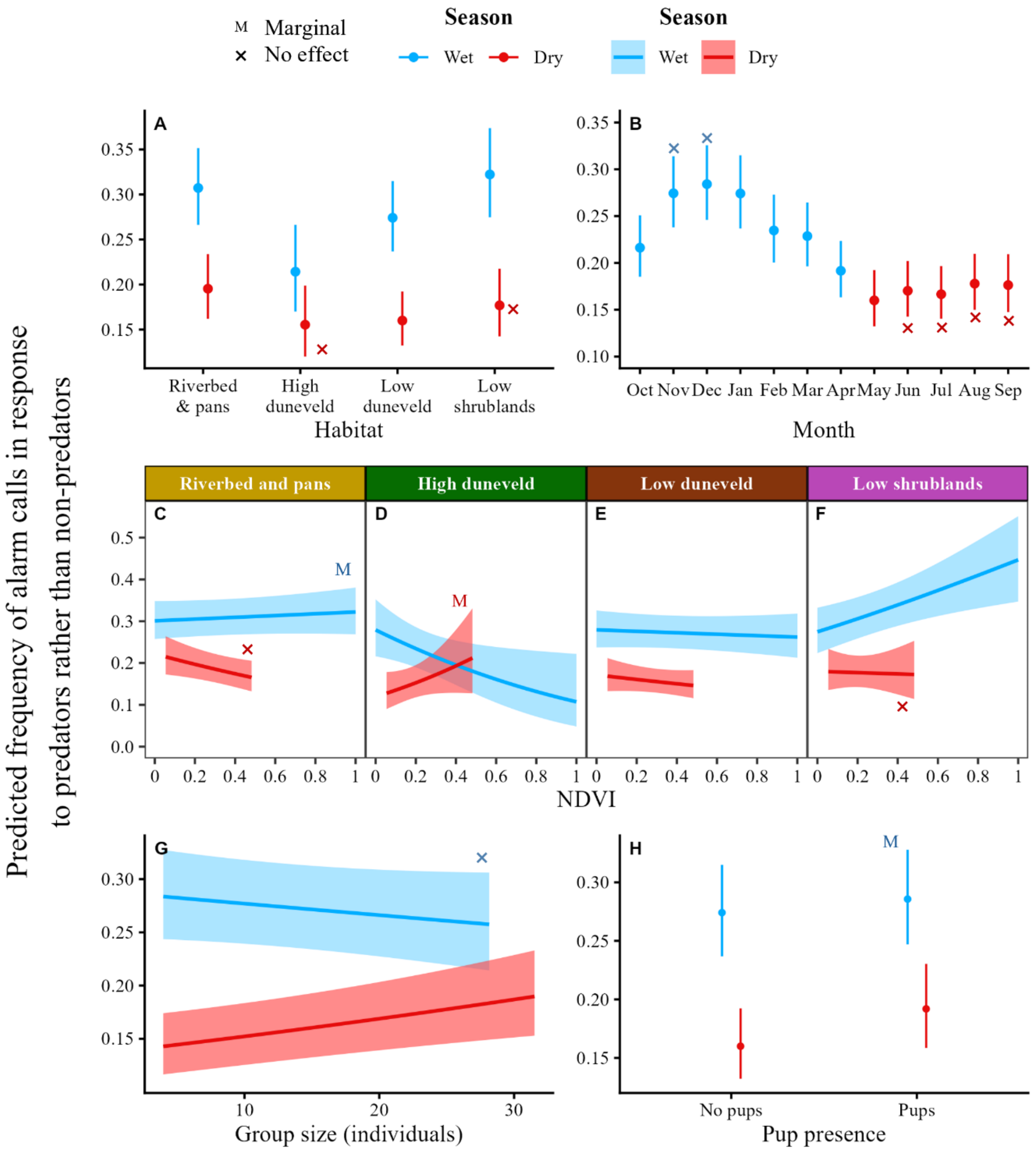

**Figure S1.10.** Predicted values for all the effects of the final model of the objective on threat discrimination **(with "Month" added as a predictor)** for the wet (blue) and dry (red) seasons for which the 95% confidence intervals did not include 0. Error bars and ribbons are the 95% confidence intervals. To observe the effect better, y axes were scaled accordingly. (A) Baseline habitat structure. **(B) Month effect.** (C, D, E, F) Habitat-specific Normalised Difference Vegetation Index (NDVI) effects predicted over a more realistic NDVI range excluding 5% of extreme values. (F) Group size effects predicted over a more realistic group size range excluding 5% of extreme values. (G) Pups.

**Table S1.27.** Estimates from threat discrimination final seasonal models including "**Fortnight**" as predictor. Confidence intervals excluding 0 are shown in bold. Abbreviations: SE, standard error; CI, confidence interval; $\sigma^2$, variance of random effects; NDVI, Normalised Difference Vegetation Index. Reference levels are low duneveld for habitat type, fine visibility, and no pups.

| | **Wet season** | | | **Dry season** | | |
|---|---|---|---|---|---|---|
| **Term** | **Estimate** | **SE** | **95% CI** | **Estimate** | **SE** | **95% CI** |
| **Fixed effects** | | | | | | |
| **Intercept** | **-1.089** | **0.118** | **-1.32 – -0.86** | **-1.79** | **0.131** | **-2.05 – -1.53** |
| *Temporal effects* | | | | | | |
| January (B) / **May (B)** | 0.214 | 0.112 | -0.01 – 0.43 | **0.222** | **0.105** | **0.02 – 0.43** |
| February (A) / **June (A)** | -0.072 | 0.118 | -0.3 – 0.16 | **0.318** | **0.107** | **0.11 – 0.53** |
| February (B) / June (B) | -0.096 | 0.117 | -0.32 – 0.13 | 0.087 | 0.111 | -0.13 – 0.3 |
| **March (A) /** July (A) | **-0.252** | **0.116** | **-0.48 – -0.03** | 0.083 | 0.112 | -0.14 – 0.3 |
| March (B) / **July (B)** | -0.012 | 0.11 | -0.23 – 0.2 | **0.26** | **0.112** | **0.04 – 0.48** |
| **April (A) / August (A)** | **-0.257** | **0.113** | **-0.48 – -0.04** | **0.274** | **0.115** | **0.05 – 0.5** |
| **April (B) / August (B)** | **-0.424** | **0.11** | **-0.64 – -0.21** | **0.238** | **0.118** | **0.01 – 0.47** |
| October (A) / Sep (A) | -0.168 | 0.112 | -0.39 – 0.05 | 0.141 | 0.125 | -0.1 – 0.39 |
| **October (B) / Sep (B)** | **-0.24** | **0.11** | **-0.46 – -0.02** | **0.353** | **0.124** | **0.11 – 0.6** |
| November (A) | 0.094 | 0.108 | -0.12 – 0.31 | | | |
| November (B) | 0.13 | 0.11 | -0.09 – 0.34 | | | |
| **December (A)** | **0.253** | **0.112** | **0.03 – 0.47** | | | |
| December (B) | 0.048 | 0.117 | -0.18 – 0.28 | | | |
| *Spatial effects* | | | | | | |
| **Riverbed and pans** | **0.161** | **0.049** | **0.06 – 0.26** | **0.245** | **0.055** | **0.14 – 0.35** |
| **High duneveld** | **-0.323** | **0.116** | **-0.55 – -0.1** | -0.037 | 0.114 | -0.26 – 0.19 |
| **Low shrublands** | **0.232** | **0.073** | **0.09 – 0.38** | 0.122 | 0.086 | -0.05 – 0.29 |
| NDVI | -0.039 | 0.04 | -0.12 – 0.04 | -0.042 | 0.046 | -0.13 – 0.05 |
| Riverbed and pans × NDVI | 0.054 | 0.044 | -0.03 – 0.14 | -0.038 | 0.05 | -0.14 – 0.06 |
| High duneveld × NDVI | -0.277 | 0.143 | -0.56 – 0 | 0.209 | 0.109 | 0 – 0.42 |
| **Low shrublands × NDVI** | **0.229** | **0.074** | **0.08 – 0.37** | 0.037 | 0.088 | -0.14 – 0.21 |
| *Social effects* | | | | | | |
| **Group size** | -0.03 | 0.026 | -0.08 – 0.02 | **0.09** | **0.028** | **0.04 – 0.14** |
| Pups | 0.057 | 0.042 | -0.02 – 0.14 | **0.227** | **0.058** | **0.11 – 0.34** |
| **Random effects** | **$\sigma^2$** | | | **$\sigma^2$** | | |
| Year | 0.105 | | | 0.131 | | |
| Group identity | 0.045 | | | 0.080 | | |

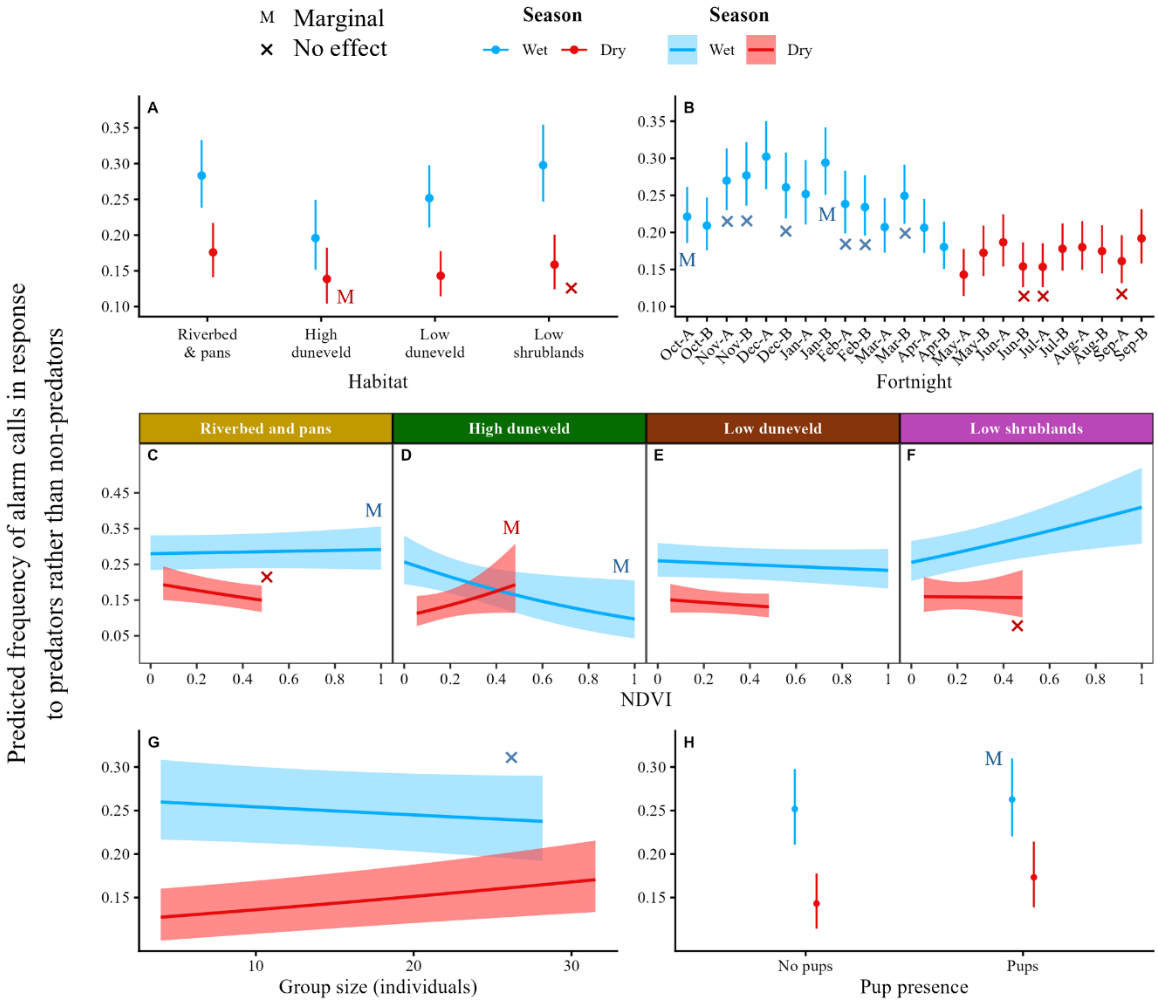

**Figure S1.11.** Predicted values for all the effects of the final model of the objective on threat discrimination **(with "Fortnight" added as a predictor)** for the wet (blue) and dry (red) seasons for which the 95% confidence intervals did not include 0. Error bars and ribbons are the 95% confidence intervals. To observe the effect better, y axes were scaled accordingly. (A) Baseline habitat structure. **(B) Fortnight effect.** (C, D, E, F) Habitat-specific Normalised Difference Vegetation Index (NDVI) effects predicted over a more realistic NDVI range excluding 5% of extreme values. (F) Group size effects predicted over a more realistic group size range excluding 5% of extreme values. (G) Pups.

**Table S1.28.** Estimates from threat discrimination final seasonal models **excluding "Riverbed and pans" as a habitat** from the dataset. Confidence intervals excluding 0 are shown in bold. Abbreviations: SE, standard error; CI, confidence interval; $\sigma^2$, variance of random effects; NDVI, Normalised Difference Vegetation Index. Reference levels are low duneveld for habitat type, fine visibility, and no pups.

| | **Wet season** | | | **Dry season** | | |
|---|---|---|---|---|---|---|
| **Term** | **Estimate** | **SE** | **95% CI** | **Estimate** | **SE** | **95% CI** |
| **Fixed effects** | | | | | | |
| **Intercept** | **-1.146** | **0.091** | **-1.33 – -0.97** | **-1.592** | **0.1** | **-1.79 – -1.4** |
| *Spatial effects* | | | | | | |
| **High duneveld** | **-0.315** | **0.116** | **-0.54 – -0.09** | -0.03 | 0.114 | -0.25 – 0.19 |
| **Low shrublands** | **0.195** | **0.074** | **0.05 – 0.34** | 0.107 | 0.086 | -0.06 – 0.28 |
| **NDVI** | **-0.066** | **0.034** | **-0.13 – 0** | -0.059 | 0.036 | -0.13 – 0.01 |
| **High duneveld × NDVI** | **-0.256** | **0.131** | **-0.51 – 0** | **0.207** | **0.097** | **0.02 – 0.4** |
| **Low shrublands × NDVI** | **0.212** | **0.067** | **0.08 – 0.34** | 0.04 | 0.079 | -0.11 – 0.19 |
| *Social effects* | | | | | | |
| Group size | 0.033 | 0.032 | -0.03 – 0.1 | 0.065 | 0.034 | 0 – 0.13 |
| **Pups** | 0.065 | 0.053 | -0.04 – 0.17 | **0.151** | **0.072** | **0.01 – 0.29** |
| **Random effects** | $\sigma^2$ | | | $\sigma^2$ | | |
| Year | 0.104 | | | 0.139 | | |
| Group identity | 0.076 | | | 0.081 | | |

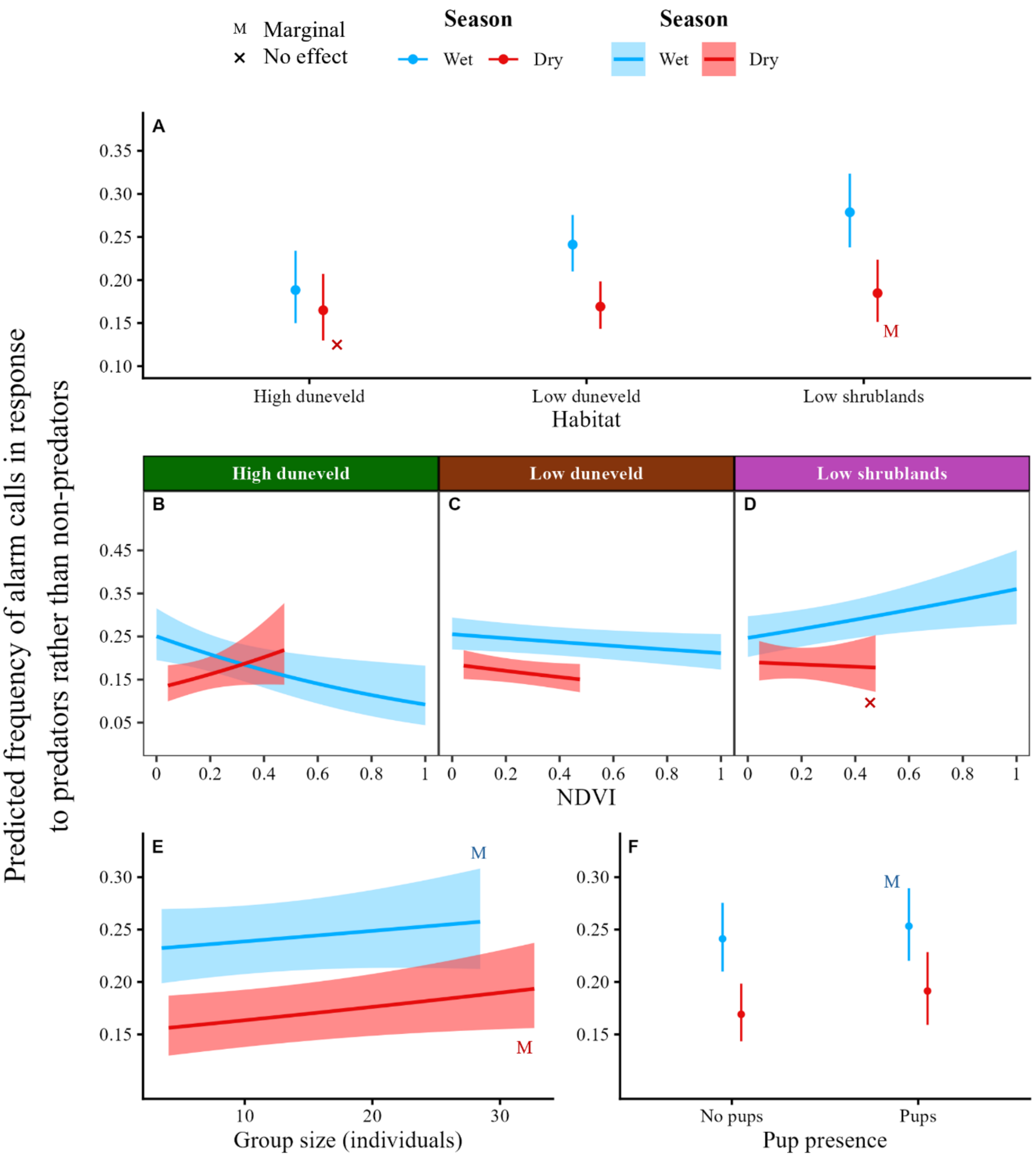

**Figure S1.12.** Predicted values for all the effects of the final model of the objective on threat discrimination **(with "Riverbed and pans" excluded from the dataset)** for the wet (blue) and dry (red) seasons for which the 95% confidence intervals did not include 0. Error bars and ribbons are the 95% confidence intervals. To observe the effect better, y axes were scaled accordingly. (A) Baseline habitat structure. (B, C, D, E) Habitat-specific Normalised Difference Vegetation Index (NDVI) effects predicted over a more realistic NDVI range excluding 5% of extreme values. (F) Group size effects predicted over a more realistic group size range excluding 5% of extreme values. (G) Pups.

#### 9. Temporal and spatial autocorrelation checks

##### *9A - Threat discrimination*

**Table S1.29.** Summary of the temporal autocorrelation test for the objective on threat discrimination. Lag-1 ACF is the correlation between residuals at time t and t+1 within a given group follow.

| Season | Mean lag-1 ACF | Median lag-1 ACF | SD lag-1 ACF | Sessions tested |
|---|---|---|---|---|
| Wet | -0.117 | -0.118 | 0.327 | 386 |
| Dry | -0.142 | -0.135 | 0.306 | 505 |

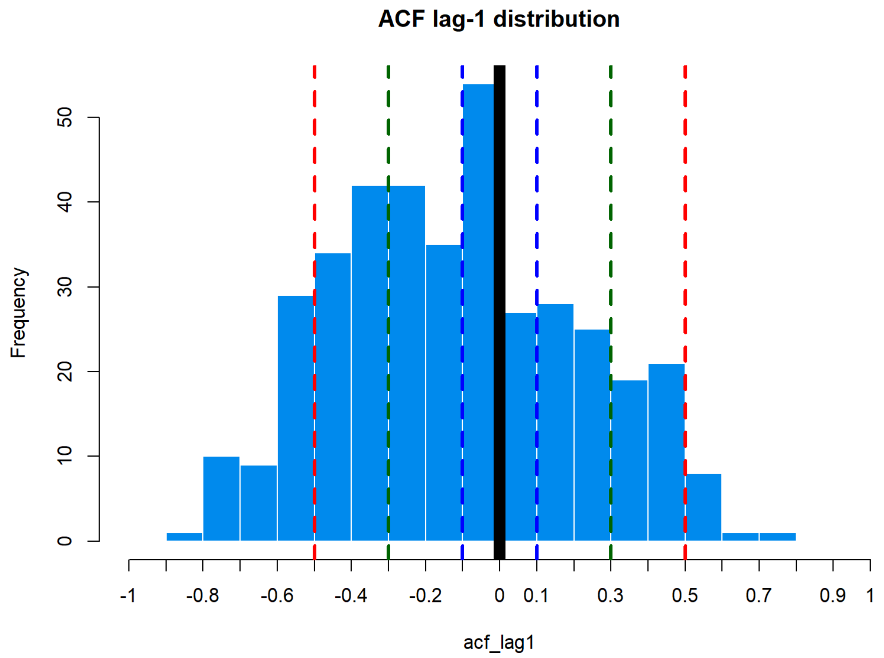

**Figure S1.13.** Wet season results from the temporal autocorrelation test. Results within the blue dashed line suggest negligible temporal autocorrelation. Results within the green dashed lines suggest weak temporal autocorrelation. Results within the red dashed lines suggest moderate temporal autocorrelation. Results beyond that suggest strong temporal autocorrelation.

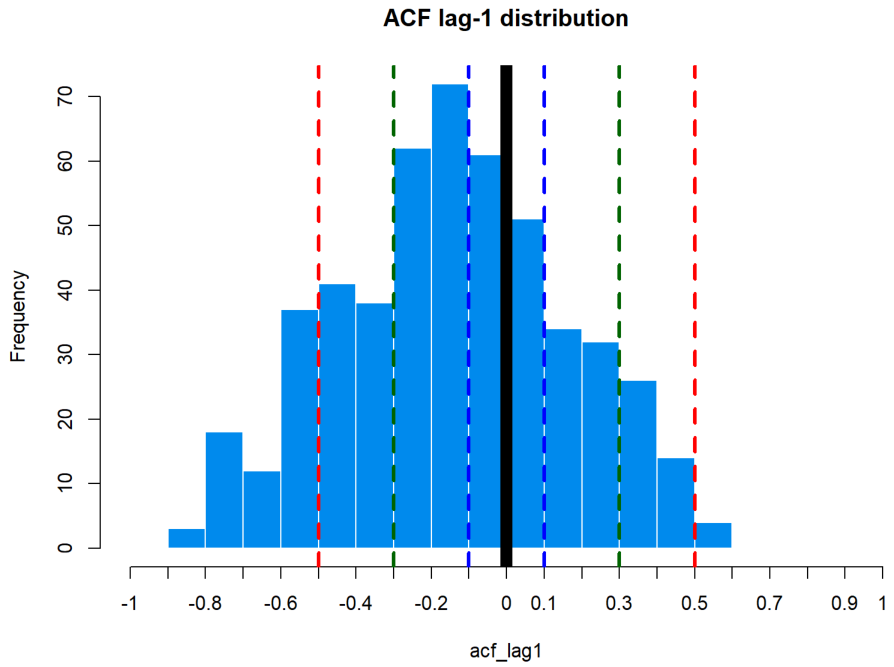

**Figure S1.14.** Dry season results from the temporal autocorrelation test. Results within the blue dashed line suggest negligible temporal autocorrelation. Results within the green dashed lines suggest weak temporal autocorrelation. Results within the red dashed lines suggest moderate temporal autocorrelation. Results beyond that suggest strong temporal autocorrelation.

Mean lag-1 ACF are mostly centred around 0, even negative, indicating negligible temporal autocorrelation.

**Table S1.30.** Spatial autocorrelation test results using Moran I test under randomisation for the threat discrimination objective.

| Season | Moran's I | P-value |
|---|---|---|
| Wet | 0.0337 | 0.0011 |
| Dry | 0.0098 | 0.1853 |

Significant in the wet season, but extremely small value for Moran's I. Therefore, spatial autocorrelation is negligible.

##### *9B - Response escalation: reaction strength*

**Table S1.31.** Summary of the temporal autocorrelation test for the objective on reaction strength. Lag-1 ACF is the correlation between residuals at time t and t+1 within a given group follow.

| Mean lag-1 ACF | Median lag-1 ACF | SD lag-1 ACF | Sessions tested |
|---|---|---|---|
| -0.176 | -0.183 | 0.307 | 560 |

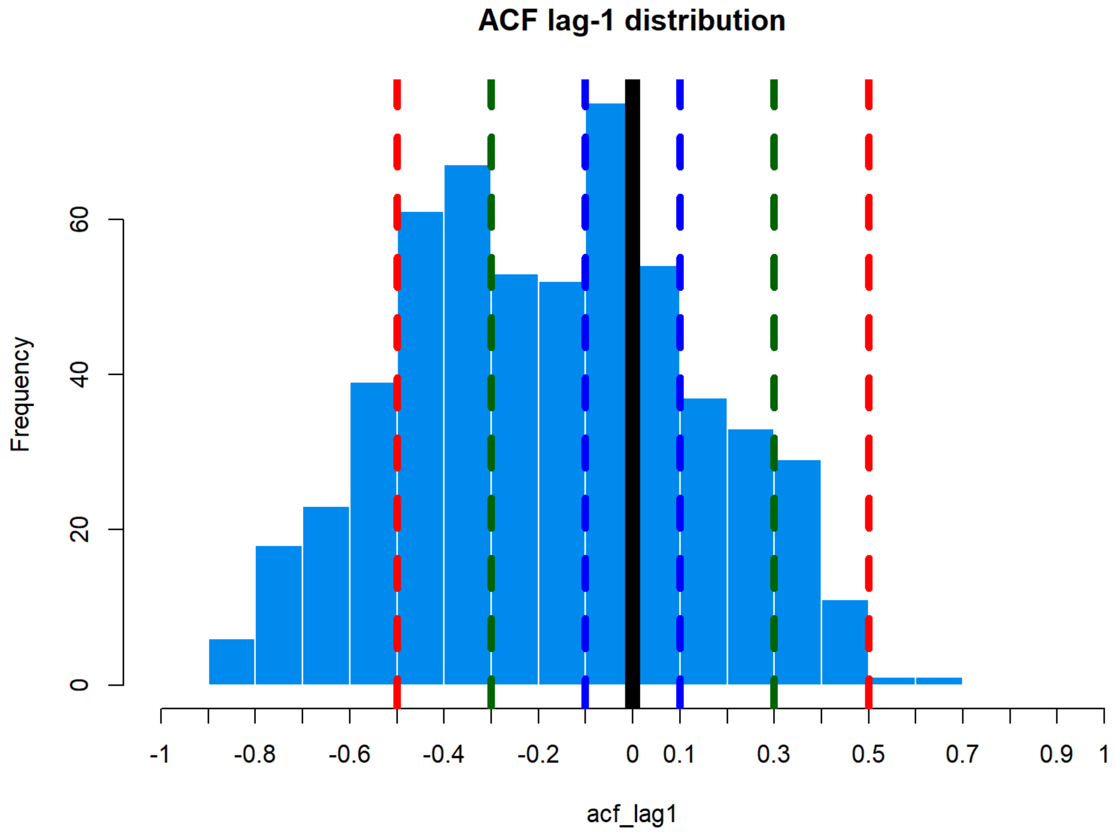

**Figure S1.15.** Results of the temporal autocorrelation test. Results within the blue dashed line suggest negligible temporal autocorrelation. Results within the green dashed lines suggest weak temporal autocorrelation. Results within the red dashed lines suggest moderate temporal autocorrelation. Results beyond that suggest strong temporal autocorrelation.

Mean lag-1 ACF is centred around 0 and slightly negative, suggesting only negligible temporal autocorrelation.

**Table S1.32.** Spatial autocorrelation test results using Moran I test under randomisation for the reaction strength objective.

| Moran's I | P-value |
|---|---|
| 0.0335 | 0.0005 |

Significant but extremely weak, therefore spatial autocorrelation is negligible.

*9C - Response escalation: reaction intensity*

**Table S1.33.** Summary of the temporal autocorrelation test for the objective on reaction intensity. Lag-1 ACF is the correlation between residuals at time t and t+1 within a given group follow.

| Mean lag-1 ACF | Median lag-1 ACF | SD lag-1 ACF | Sessions tested |
|---|---|---|---|
| -0.175 | -0.200 | 0.310 | 313 |

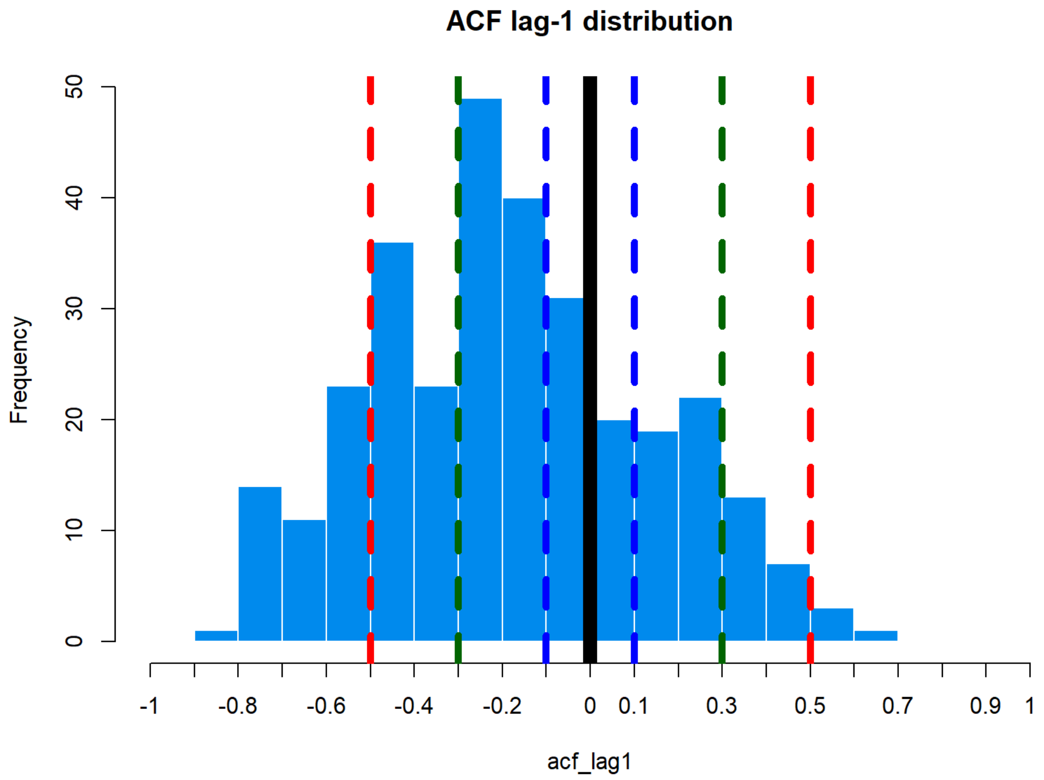

**Figure S1.16.** Results of the temporal autocorrelation test. Results within the blue dashed line suggest negligible temporal autocorrelation. Results within the green dashed lines suggest weak temporal autocorrelation. Results within the red dashed lines suggest moderate temporal autocorrelation. Results beyond that suggest strong temporal autocorrelation.

Mean lag-1 ACF is centred around 0, suggesting negligible temporal autocorrelation.

**Table S1.34.** Spatial autocorrelation test results using Moran I test under randomisation for the reaction intensity objective.

| Moran's I | P-value |
|---|---|
| 0.0337 | 0.0006 |

Slight significance, but very weak Moran's I value, so spatial autocorrelation can only be negligible.
